# Multi-timescale learning signals in *Drosophila* dopaminergic neurons

**DOI:** 10.64898/2026.06.08.729969

**Authors:** Woochan Choi, Shyam Srinivasan, Dhruv Grover

## Abstract

Learning often requires inference over hidden task structure, including features that predict outcomes. In mammals, prediction-error signaling by midbrain dopamine neurons is considered central to learning, but how these neurons reflect the progression and stability of learning remains unclear. Using *Drosophila* to monitor calcium activity at trial-by-trial resolution during aversive conditioning, we found that PPM3 dopaminergic neurons exhibit hallmark prediction-error responses in which activity shifted from the unconditioned stimulus (US) to the conditioned stimulus (CS), was suppressed when an expected US was omitted, and increased when the US exceeded expectations. Strikingly, the same neurons also exhibited slower state-like dynamics across trials, including tonic activity transitions that emerged with learning and tracked the acquisition of learned behavior, and perturbation-related dynamics that were briefly disrupted when expectations were violated. Increasing task demands by inserting a temporal gap between the CS and US (trace conditioning) delayed both response types to later trials, accompanied by corresponding delays in behavior acquisition. In addition, dopaminergic activity developed an anticipatory response that tracked the expected timing of the US during the gap. These findings reveal that a single dopaminergic neuron type integrates moment-to-moment prediction errors, expected outcome timing, and a longer-timescale signal reflecting learning stabilization, establishing *Drosophila* as a bona fide model for dissecting the neural mechanisms that shape learning under changing task demands.

---

The core of modern reinforcement learning theory is the Temporal Difference (TD) model, a computational framework in which an agent learns to predict future outcomes by continuously updating its value estimates based on the discrepancy between expected and actual inputs^1,2^. This discrepancy, termed the Reward Prediction Error (RPE), serves as a teaching signal that drives learning by updating expectations about future outcomes. A major advance in neuroscience came with the proposal that midbrain dopamine neurons may instantiate this computation in the brain, broadcasting RPE signals that guide adaptive behavior^3–5^. Consistent with this framework, phasic dopamine activity increases when an outcome is better than expected, remains near baseline when an outcome is fully predicted, and decreases when an expected outcome is omitted or worse than predicted^3,6,7^.

Although classical reinforcement learning theories describe dopamine as reporting a scalar prediction error that updates value estimates based on external cues and outcomes, adaptive learning in natural environments cannot rely on observable events alone. Animals must also infer the hidden structure of a task, including whether observations arise from the same context, whether contingencies remain stable, and which past events are causally responsible for current outcomes^8–12^. For example, when an expected reward suddenly fails to appear, an animal must determine whether this reflects random variability, a change in context, or a shift in the underlying rules governing outcomes. Learning therefore depends not only on experienced outcomes, but also on an internal belief about latent task states that is continuously updated as evidence accumulates. In line with this view, recent theoretical and experimental work suggests that dopaminergic prediction-error signals can be shaped by hidden-state inference across time rather than by a fixed sequence of observable events alone^13–16^. Yet, despite the central importance of hidden learning states in modern learning theory, how such states are represented in neural activity during learning remains unclear. Identifying trial-resolved neural signals of learning state may therefore be essential for understanding how internal predictions emerge, stabilize, and reorganize across experience.

In the *Drosophila* brain, dopaminergic neurons (DANs) projecting to the Mushroom Body have been shown to provide the reinforcement signals that drive associative olfactory learning, instructing synaptic plasticity between the Kenyon cells and Mushroom Body output neurons to assign positive or negative value to sensory cues^17–20^. This reinforcement framework has successfully explained much of the fly’s behavioral plasticity^21^, positioning DANs as teaching signals that reinforce or weaken learned associations^18,20,22,23^. However, whether these neurons carry richer learning-related computations remains largely unknown. Although computational models have proposed mechanisms by which Mushroom-Body-projecting DANs could encode prediction errors and learning-dependent signals^24–26^, direct physiological evidence for prediction error-like dynamics in fly dopamine neurons has remained elusive. As a result, DANs have been interpreted primarily as conveyors of external reinforcement, rather than as neural signals shaped by internal expectations or latent learning states.

Here, we build on prior work demonstrating that the PPM3 DANs that project to the Ellipsoid Body substructure of the Central Complex (hereafter referred to as PPM3-EB), are required for visual aversive delay and trace conditioning in *Drosophila*^27^, to examine how these neurons encode learning over time. Using *in vivo* calcium imaging during trial-by-trial aversive conditioning, we find that PPM3-EB neurons exhibit hallmark prediction-error activity shifts from the punishment to the predictive visual cue during learning, remains near baseline for expected outcomes, decreases when an expected punishment is omitted, and increases when the punishment exceeded expectations. Beyond these rapid, stimulus-locked signals, we also observed a slower evolution of activity across trials that tracks the progression of learning and its stability over time. This slower component is transiently disrupted when expectations are violated, suggesting sensitivity to changes in the inferred task structure. Increasing task demands through trace conditioning delays the emergence of both rapid prediction-error responses and slower learning-related dynamics, with corresponding delays in behavioral acquisition, while also revealing an anticipatory signal that tracks the expected timing of the punishment across the interval between the predictive visual cue and punishment. Together, these findings suggest that PPM3-EB neurons integrate prediction-error signaling, temporal expectation, and slower learning-dependent changes in internal state across multiple timescales during associative learning.

## Results

### PPM3-EB dopaminergic neurons exhibit prediction-error-like responses

To examine how PPM3-EB dopaminergic neurons encode associative learning, we performed *in vivo* two-photon calcium imaging of these neurons^27,28^ while the flies underwent a visual delay conditioning task (Fig. 1a, b). In this paradigm, an upright “T”-shaped visual cue served as the conditioned stimulus (CS), paired with a heat punishment delivered via an infrared laser that raised the body temperature to 35 °C as the unconditioned stimulus (US), overlapping temporally with the CS (Fig. 1a, c).

**Figure 1.**
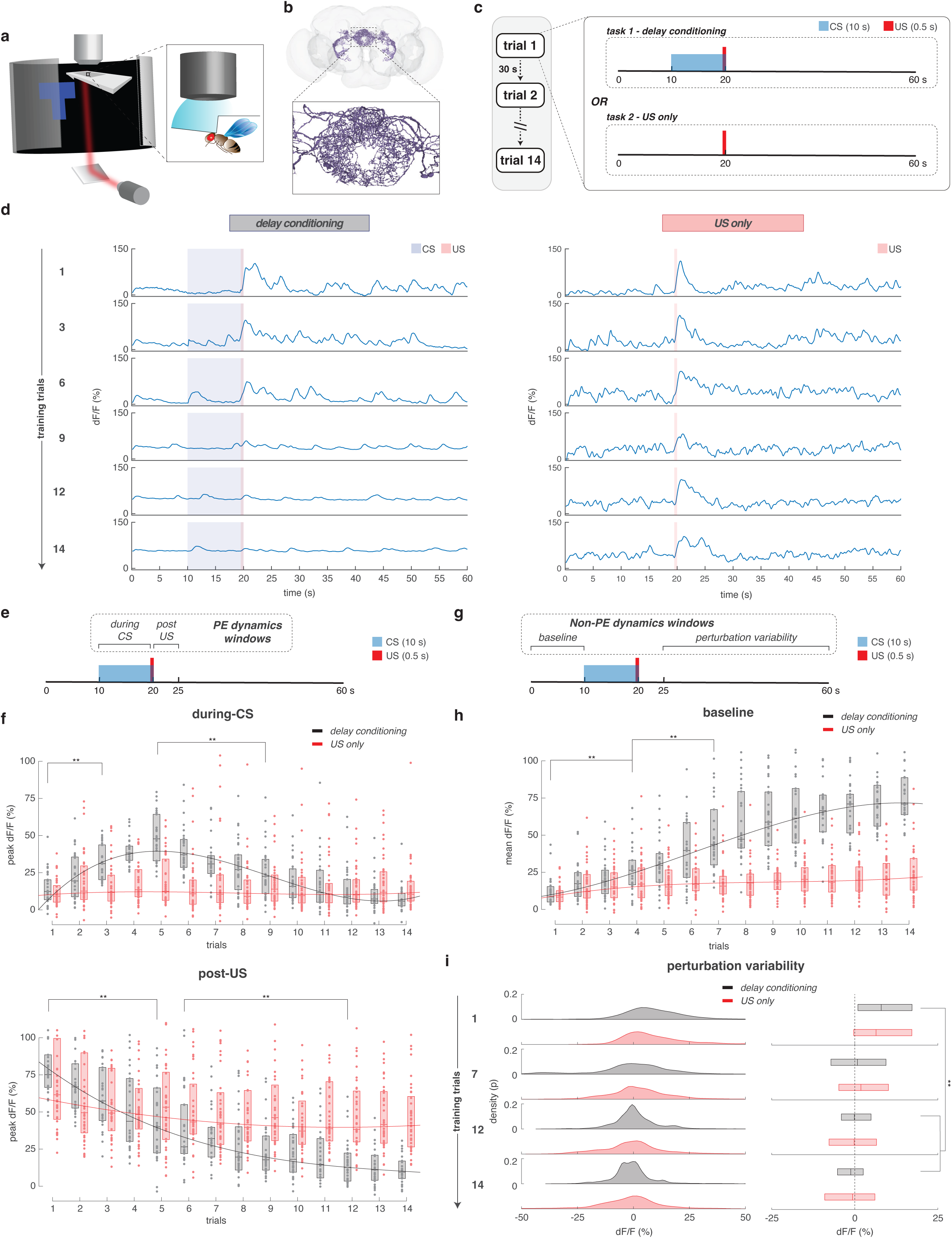
Ratiometric calcium imaging of EB-projecting PPM3 dopaminergic neurons during delay conditioning show phasic prediction-error-like responses and tonic learning-dependent activity. **(a)** Visual conditioning assay coupled with two-photon *in vivo* brain imaging - tethered fly under an objective (inset) shown an upright-T paired with heat. **(b)** 3D visualization of dopaminergic PPM3-EB (ExR2) neuron cluster. This panel is created using the FlyWire anatomical reconstruction. Inset: zoomed-in display of synaptic contact sites in the EB forming a ring structure. **(c)** Flies were tested under either delay conditioning wherein a CS (either an upright- or inverted-T) was paired with US (heat) or a US-only protocol wherein the US was presented without CS for 14 trials. **(d)** Ratiometric calcium imaging of a *c346-Gal4>>UAS-GCaMP7f;UAS-myr-tdTomato* female during delay conditioning (left) and US-only (right). Shown, dF_ratio_/F_ratio_ activity (trials 1, 3, 6, 9, 12, and 14). **(e)** Time windows used for determining prediction-error-related statistics during delay conditioning and matched controls. For prediction error-specific dynamics, activity during CS (10-19.5 s) and post-US (20-25 s) were considered. **(f)** Peak dF_ratio_/F_ratio_ activity during CS (10-19.5 s, top), and post-US (20-25s, bottom) for delay conditioning (black, n = 42 flies) and US-only (red, n = 41 flies). Shown on top, quadratic (second degree) polynomial curve fits, and bottom, single-term exponential curve-fits through median dF_ratio_/F_ratio_ activity. **(g)** Time windows used for determining non-prediction error-related statistics during delay conditioning and matched controls. For non-prediction error-related dynamics, the (pre-CS) baseline activity (0-10 s) and perturbation variability after the post-US window (25-60 s) were considered. **(h)** Mean dF_ratio_/F_ratio_ activity during the pre-CS baseline period (0-10 s) for delay conditioning (black) and US-only (red). Shown are quadratic (second degree) polynomial curve fits through median dF_ratio_/F_ratio_ activity. **(i)** Distributions of perturbation variability (relative to pre-CS baseline mean) 5 s after US (from 25-60 s) for delay conditioning (black) and US-only (red). Shown are kernel probability density estimate (left) and boxplot (right) representations for trials 1, 7, 12, 14. Equality of group variances was tested using Brown-Forsythe variant ANOVA with Holm multiplicity correction. Boxplot center (median), and edges (IQR). Scatters represent single-fly metrics. Groups compared using two-factor ART-ANOVA. n.s. indicates not significant, * is p-value < 0.05, ** p-value < 0.01.

We first examined neural activity time-locked to task events across training. Early in conditioning, PPM3-EB neurons exhibited strong phasic calcium responses to the unexpected punishment. Across successive trials, these US-evoked responses progressively diminished as the punishment became predicted by the CS (Fig. 1d–f, Fig. S1a–b). In contrast, in a US-only condition in which punishment was delivered without the predictive visual cue, US-evoked responses remained stable across trials, indicating that the attenuation of activity depends on learned prediction rather than sensory adaptation.

In parallel with the reduction in punishment-evoked responses, PPM3-EB activity shifted during learning to become increasingly aligned with the predictive visual cue. Phasic calcium transients that were initially time-locked to the punishment progressively emerged earlier in the trial and became associated with CS presentation, accompanied by a reduction in US-locked activity (Fig. 1f). This progressive shift from outcome-locked to cue-locked activity is consistent with prediction-dependent updating during associative learning.

Together, these dynamics demonstrate that PPM3-EB neurons exhibit hallmark features of event-locked prediction-error-like signaling, including sensitivity to outcome expectancy and a learning-related shift in activity from punishment to predictive cue.

### PPM3-EB dopaminergic neurons exhibit slow trial-by-trial activity dynamics

To determine whether PPM3-EB activity evolves across learning beyond event-locked responses, we analyzed calcium dynamics across individual training trials independent of stimulus timing. This revealed a second component of neural activity that was not strictly time-locked to either the predictive visual cue or the punishment.

First, PPM3-EB neurons exhibited a sustained increase in baseline calcium activity during delay conditioning that emerged progressively across training (Fig. 1g–h). This tonic elevation was absent in the US-only condition, where repeated punishment alone failed to produce a comparable increase in baseline activity. Likewise, no increase was observed in the CS-only condition (Fig. S2a–c), indicating that the elevated baseline activity depends on the association between cue and outcome rather than sensory stimulation or punishment exposure alone.

Second, we quantified trial-to-trial fluctuations in calcium activity during delay conditioning, revealing a systematic reduction in variability across training (Fig. 1i). In contrast, variability remained elevated in the US-only condition despite repeated punishment exposure, suggesting that stabilization of neural activity depends on predictive structure rather than aversive stimulation alone. Notably, variability increased immediately following the first punishment exposure in both conditioning and US-only conditions (Fig. S3a–b), indicating that punishment transiently increases neural fluctuations before activity stabilizes over training.

We refer to these two slow components as a tonic baseline shift, reflecting sustained elevation in baseline activity across trials, and perturbation variability, reflecting trial-to-trial fluctuations in calcium activity. Together, these findings identify a slower timescale of PPM3-EB activity that evolves across training and is distinct from event-locked responses, characterized by a progressive increase in baseline activity and reduced trial-to-trial variability during associative conditioning.

### Prediction error responses and perturbation variability exhibit dissociable coding during rule violations

To further distinguish fast prediction-error responses from slower trial-by-trial dynamics, we examined PPM3-EB activity during violations of learned contingencies following conditioning^3,5^. After training, expected punishment was either omitted or delivered at a greater intensity, allowing us to probe neural responses to both negative and positive deviations from learned predictions (Fig. 2a).

**Figure 2.**
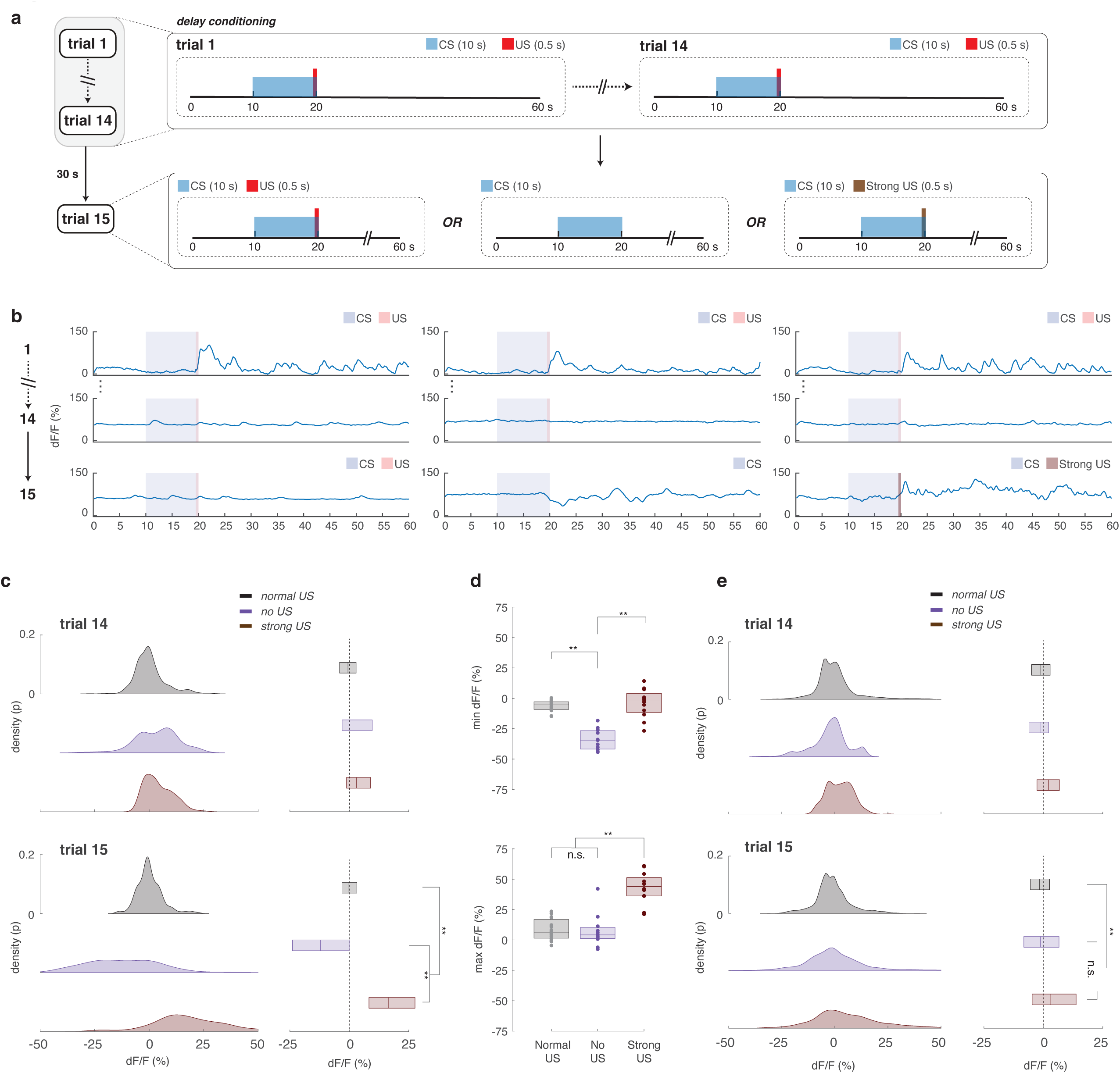
Delay conditioning outcome violations reveal signed prediction errors and re-engaged perturbation variability in EB-PPM3 calcium dynamics. **(a)** Illustration of the experimental design used for testing dopamine prediction error signaling, 14 trials of typical delay conditioning task, followed by a probe trial 15 that tests one of three conditions – typical delay conditioning (left), omission of US (CS-only presentation, middle), or delay conditioning with a stronger-US that delivers a heat punishment raising the fly body temperature to 40 °C instead of the typical 35 °C (right). **(b)** Ratiometric calcium imaging of a *c346-Gal4>>UAS-GCaMP7f;UAS-myr-tdTomato* females for each of the three learning conditions in **(a)**, during typical delay conditioning (left), 14 trials of delay conditioning followed by omission of US in trial 15 (middle), and 14 trials of delay conditioning followed by stronger-US in trial 15 (right). Shown, dF_ratio_/F_ratio_ activity (trials 1, 14, and probe trial 15). **(c)** Comparison of the distributions of activity (relative to pre-CS baseline mean) during the post-US interval (20-25 s) for trial 14 (top row) and probe trial 15 (bottom row) for the three conditions in **(a, b)**. Shown are kernel probability density estimate (left column) and boxplot (right column) representations. **(d)** Minimum (top) and maximum (bottom) dF_ratio_/F_ratio_ activity (relative to pre-CS baseline mean) during post-US interval (20-25 s) in probe trial 15 for **(a, b)**. **(e)** Comparison of the distributions of perturbation variability (relative to pre-CS baseline mean) 5 s after US (25-60 s) for trial 14 (top row) and probe trial 15 (bottom row) for the three conditions in **(a, b)**. Shown are kernel probability density estimate (left column) and boxplot (right column) representations. Equality of group variances was tested using Brown-Forsythe variant ANOVA with Holm multiplicity correction. In **(c-e)**, shown are data from typical delay conditioning (black, n = 19 flies), omission of US condition (purple, 11 flies), and stronger-US condition (brown, n = 12 flies). Boxplot center (median), and edges (IQR). Group distributions compared using Kruskall-Wallis and post-hoc bonferroni-corrected unpaired two-sided Mann–Whitney U tests. n.s. indicates not significant, * is p-value < 0.05, ** p-value < 0.01.

Consistent with temporal-difference models of prediction-error signaling^3,5,29^, PPM3-EB neurons exhibited bidirectional, signed responses to outcome violations. Omission of an expected punishment produced a transient suppression of neural activity below baseline, consistent with a negative prediction error. Conversely, increasing punishment intensity (raising the body temperature to 40 °C instead of 35 °C) at the expected time elicited an enhanced phasic response, consistent with a positive prediction error (Fig. 2b–d). These effects were absent in the US-only controls (Fig. S4), indicating that they depend on learned expectations rather than stimulus properties alone.

We next examined perturbation variability, the slower trial-to-trial fluctuation in calcium activity identified in Fig. 1i. In contrast to prediction-error responses, perturbation variability did not depend on the direction of the outcome violation. Instead, both punishment omission and increased punishment elicited a transient increase in variability relative to baseline, regardless of whether the violation was better or worse than expected (Fig. 2e). Thus, prediction-error responses encoded signed expectation violations, whereas perturbation variability increased irrespective of violation direction.

Together, these results demonstrate that fast event-locked prediction-error responses and slower perturbation variability represent dissociable components of PPM3-EB signaling, differing in both timescale and computational coding properties.

### Slow dopaminergic activity exhibits state transitions distinct from prediction-error dynamics

The analyses above suggested that perturbation variability and tonic baseline activity evolve across training in a manner distinct from event-locked prediction-error responses. We therefore asked whether these slower components follow a different temporal update structure than phasic activity.

To address this, we compared models describing trial-by-trial dynamics of fast and slow components of PPM-EB activity. Because the timing of neural transitions varied across individuals, simple trial averaging can obscure abrupt changes in activity structure^30,31^. We therefore first applied change-point analysis (Methods, Change-point analysis) to identify potential discontinuities in neural activity across training (Fig. 3a–c).

**Figure 3.**
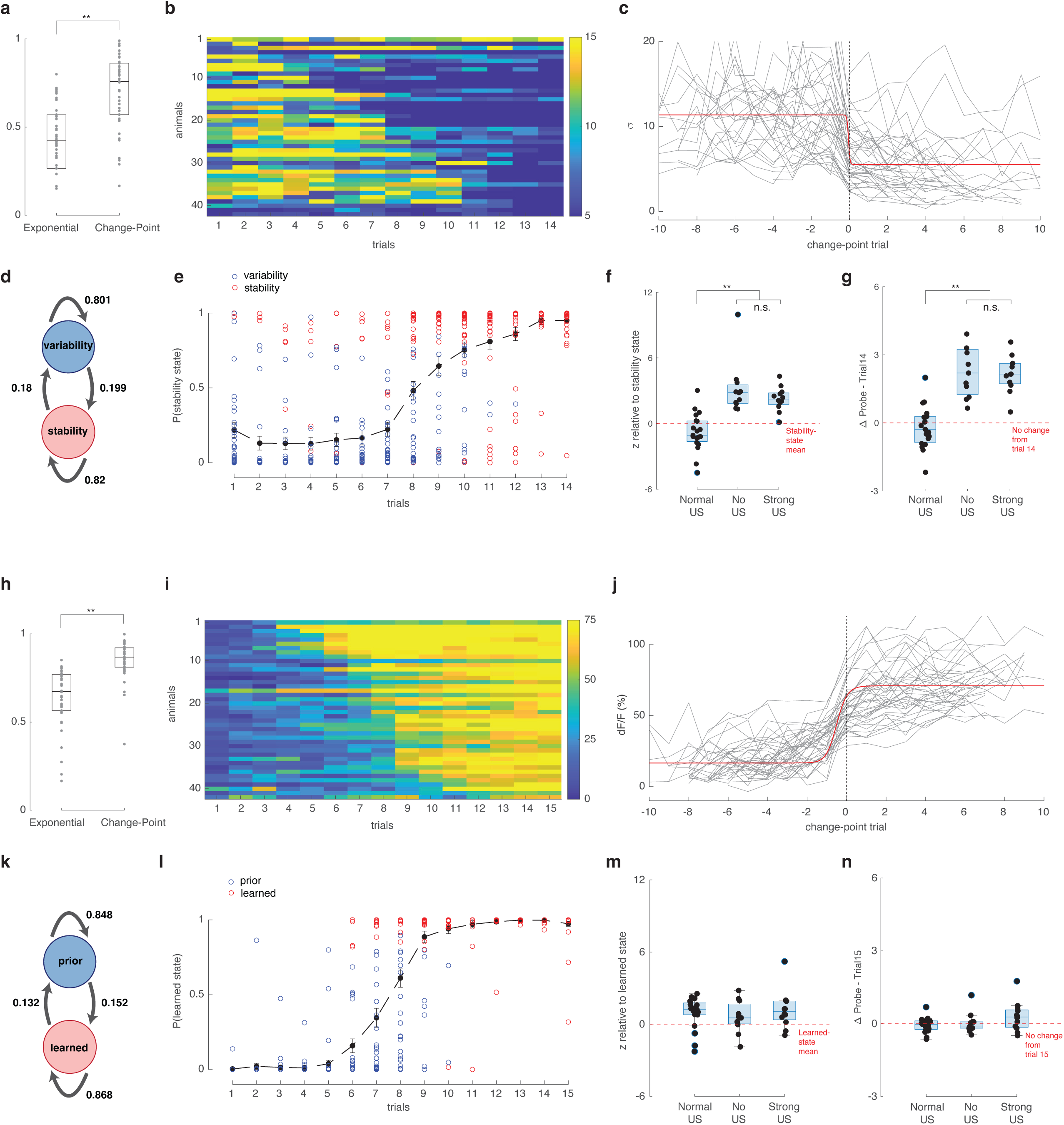
EB-PPM3 calcium dynamics reveal prediction-error responses decay gradually, whereas tonic activity shifts abruptly with state transitions. **(a)** R-squared goodness of fit of exponential and change-point models applied to per-trial standard deviations of dF_ratio_/F_ratio_ perturbation variability (relative to pre-CS baseline mean) 5 s after US (25-60 s) during delay conditioning (n = 42 flies). **(b)** Heatmap of modeled data from **(a)** in ascending order of each animal’s change point trial that partitions the data into two regions, minimizing the sum of the residual (squared) error of each region from its local mean. **(c)** Same data as **(b)**, trial aligned to center (trial 0) on each animal’s respective change point trial (gray lines). Red curve indicates sigmoidal curve fit to the per-trial median of the change point trial-aligned data. **(d)** A two-state hidden Markov model was fit to the per-trial standard deviations of perturbation variability in **(b)**, shown are the transition probabilities of estimated latent variability and stability states during the learning process. **(e)** Inferred probability of being in the variability (blue) or stability (red) state for each trial across animals. **(f)** Probe-trial (trial 15) responses were evaluated relative to the stability-state distribution for each of the three learning conditions in (Fig. 2c). Normalized deviations of trial 15 values from the stability-state were determined using z-scores with respect to the stability-state mean. **(g)** Change in the normalized responses from last learning trial (trial 14) to the probe trial (trial 15) for each of the three learning conditions in **(2c)**. Δ ∼ 0 indicates no change and stability state holds, Δ > 0 indicates increased activity, Δ < 0 indicates decrease in activity. **(h)** R-squared goodness of fit of exponential and change-point models applied to per-trial mean pre-CS baseline dF_ratio_/F_ratio_ activity (0-10 s) during delay conditioning (n = 42 flies). **(i)** Heatmap of modeled data from **(h)** in ascending order of each animal’s change point trial that partitions the data into two regions, minimizing the sum of the residual (squared) error of each region from its local mean. **(j)** Same data as **(i)**, trial aligned to center (trial 0) on each animal’s respective change point trial (gray lines). Red curve indicates sigmoidal curve fit to the per-trial median of the change point trial-aligned data. **(k)** A two-state hidden Markov model was fit to the per-trial mean baseline activity level in **(i)**, shown are the transition probabilities of estimated latent prior and learned states during the learning process. **(l)** Inferred probability of being in the prior (blue) or learned (red) state for each trial across animals. **(m)** Probe-trial (trial 16) responses were evaluated relative to the learned-state distribution for each of the three learning conditions in (fig. 2c). Normalized deviations of trial 16 values from the learned-state were determined using z-scores with respect to the learned-state mean. **(n)** Change in the normalized responses from last learning trial (trial 15) to the probe trial (trial 16) for each of the three learning conditions in **(2c)**. Δ ∼ 0 indicates no change and stable learned state holds, Δ > 0 indicates increased activity, Δ < 0 indicates decrease in activity. Multiple group comparisons performed using Kruskall-Wallis and post-hoc bonferroni-corrected unpaired two-sided Mann–Whitney U tests. Two group comparisons performed using unpaired two-sided Mann–Whitney U tests. n.s. indicates not significant, * is p-value < 0.05, ** p-value < 0.01.

This analysis revealed that slow components of PPM3-EB activity were better described by step-like transitions than by gradual decay^3,32^. Specifically, both perturbation variability and tonic baseline activity exhibited abrupt shifts between distinct activity regimes^9,13,14^. In contrast, event-locked prediction-error responses were better captured by gradual, exponential-like changes across trials, consistent with continuous updating rather than discrete transitions (Fig. S5a–c). These findings indicate that slow and fast components of PPM3-EB activity follow fundamentally different temporal update profiles.

We next asked whether these transitions could be formally captured as discrete activity states^8,11,33^. To do so, we applied a hidden Markov model (HMM), which infers latent activity states and probabilistic transitions from trial-resolved signals without imposing a fixed transition time^34–36^. HMM analysis revealed that perturbation variability transitioned from a high-variability state early in training to a lower-variability state after learning stabilized (Fig. 3d–e). Similarly, tonic baseline activity transitioned from a low-activity state to a higher-activity state over training (Fig. 3k–l). In both cases, inferred state probabilities remained stable within epochs and shifted sharply near the transition points, consistent with abrupt transitions in neural activity organization. Importantly, these transitions emerged despite constant task contingencies, indicating that PPM3-EB activity reorganizes across learning even within a fixed cue-outcome structure.

We next examined how these inferred states respond to transient violations of learned expectations. Perturbation variability remained sensitive to outcome violations, transiently deviating from the low-variability state following both punishment omission and increased punishment (Fig. 3f–g). In contrast, tonic baseline activity remained largely stable during these transient violations (Fig. 3m–n). This dissociation suggests that perturbation variability reflects a flexible updating-related component of the slow neural dynamics, whereas tonic baseline activity reflects a more stable learned state.

Together, these findings indicate that slow PPM3-EB activity is better described by state dynamics with abrupt transitions, whereas prediction-error responses follow continuous update rules. Thus, dopaminergic signaling in PPM3-EB neurons is organized across fundamentally different temporal update regimes.

### Task structure jointly delays prediction-error and learning-state dynamics

We next asked whether the temporal structure of the conditioning modulates both fast prediction-error responses and slower learning-state dynamics in PPM3-EB neurons. To address this, we compared delay conditioning with trace conditioning, in which a temporal gap separates the predictive visual cue from the punishment, increasing temporal credit-assignment demands (Fig. 4a-b).

**Figure 4.**
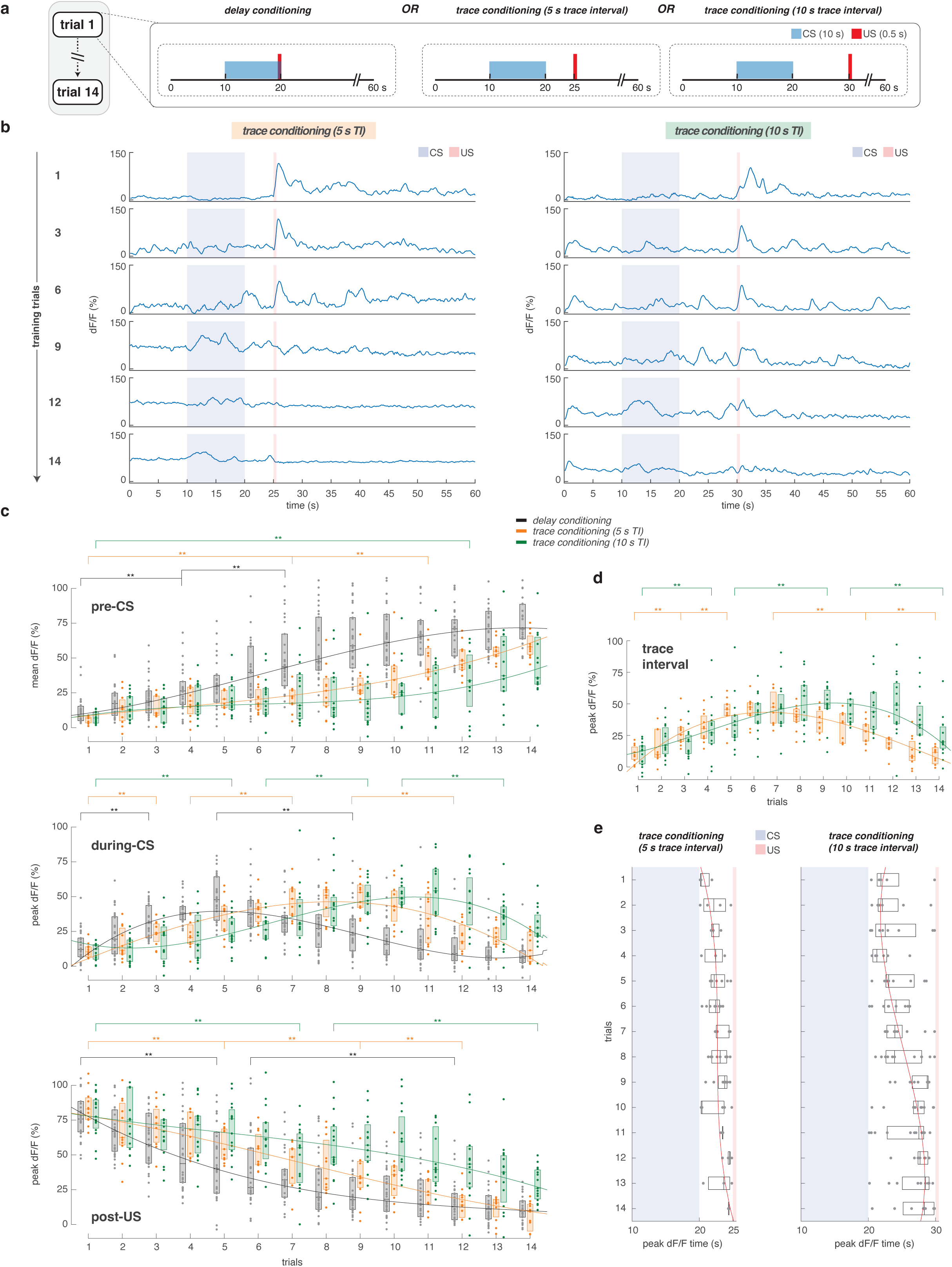
Trace conditioning delays multiple PPM3 dopaminergic calcium activity components. **(a)** Illustration of the two forms of associative learning – delay and trace conditioning. Flies were conditioned for 14 trials under either delay conditioning where the CS and US overlap (left), or trace conditioning, where CS and US are separated by a temporal gap (trace interval) of 5 s (middle) or 10 s (right). **(b)** Ratiometric calcium imaging of a *c346-Gal4>>UAS-GCaMP7f;UAS-myr-tdTomato* females during trace conditioning with a 5 s trace interval (left) and 10 s trace interval (right). Shown, dF_ratio_/F_ratio_ activity (trials 1, 3, 6, 9, 12, and 14). **(c)** Top row, mean dF_ratio_/F_ratio_ activity during the pre-CS baseline period (0-10 s) for delay conditioning (black, n = 42 flies), and trace conditioning with trace intervals of 5s (yellow, n = 11 flies), and 10 s (green, n = 15 flies). Middle and bottom, peak dF_ratio_/F_ratio_ activity during CS, and 5 s interval post-US for delay conditioning (black), and trace conditioning with trace intervals of 5 s (yellow), and 10 s (green) respectively. **(d)** Peak dF_ratio_/F_ratio_ activity during the trace interval of trace conditioning with intervals of 5 s (yellow), and 10 s (green). **(e)** Timing of the highest local maxima peak of dF_ratio_/F_ratio_ activity over trials during the trace interval of trace conditioning (left, 5 s interval; right, 10 s interval). In **(c-e)**, shown are quadratic (second degree) polynomial curve fits through median dF_ratio_/F_ratio_ activity. Boxplot center (median), and edges (IQR). Scatters represent single-fly metrics. Groups compared using two-factor ART-ANOVA. n.s. indicates not significant, * is p-value < 0.05, ** p-value < 0.01.

Across conditions, both fast event-locked responses and slower trial-by-trial dynamics were systematically delayed under trace conditioning (Fig. 4c-d). Specifically, the CS-aligned and US-evoked prediction-error responses emerged later across training when a temporal gap separated the visual cue from the punishment. This delay was accompanied by corresponding shifts in suppression and enhancement of punishment-evoked responses, consistent with slower formation of predictive associations under the increased cognitive demands imposed by the introduction of a trace interval in trace conditioning.

Strikingly, slow components exhibited a parallel delay. The emergence of tonic baseline activity and the transition in perturbation variability were both shifted later under trace conditioning relative to delay conditioning, and the transition timing scaled with the CS–US interval (Fig. 4c–d, S6–S7). Thus, increasing temporal separation between the CS and US delays not only fast prediction-error dynamics but also the slower reorganization of neural activity across training.

In addition, trace conditioning revealed a progressive shift in peak PPM3-EB activity toward the expected time of punishment during the CS-US interval (Fig. 4g). This anticipatory activity emerged gradually across training and scaled with the imposed temporal delay, suggesting that PPM3-EB activity reflects a learned estimate of expected punishment timing, consistent with previous studies implicating dopamine in interval timing and temporal representation^37,38^.

Together, these findings demonstrate that task structure jointly shapes fast prediction-error responses and slower learning-state dynamics, indicating that dopaminergic activity is organized across multiple learning timescales.

### Tonic state transitions track learned behavior

We next asked whether tonic state transitions in PPM3-EB neurons correspond to behavioral acquisition of learned avoidance. Because individual flies exhibited variability in the timing of transition from low to high tonic activity, we quantified the distribution of tonic baseline change-point trials across animals under different task structures.

During delay conditioning, most animals reached the tonic transition within the first half of training, with the average transition occurring around mid-training. A similar transition profile was observed during short (5 s trace interval) trace conditioning, although transitions were modestly delayed relative to delay conditioning. In contrast, long (10 s trace interval) trace conditioning produced a pronounced rightward shift in transition timing, with many animals failing to reach the tonic state transition within the training window (Fig. 5a–b).

**Figure 5.**
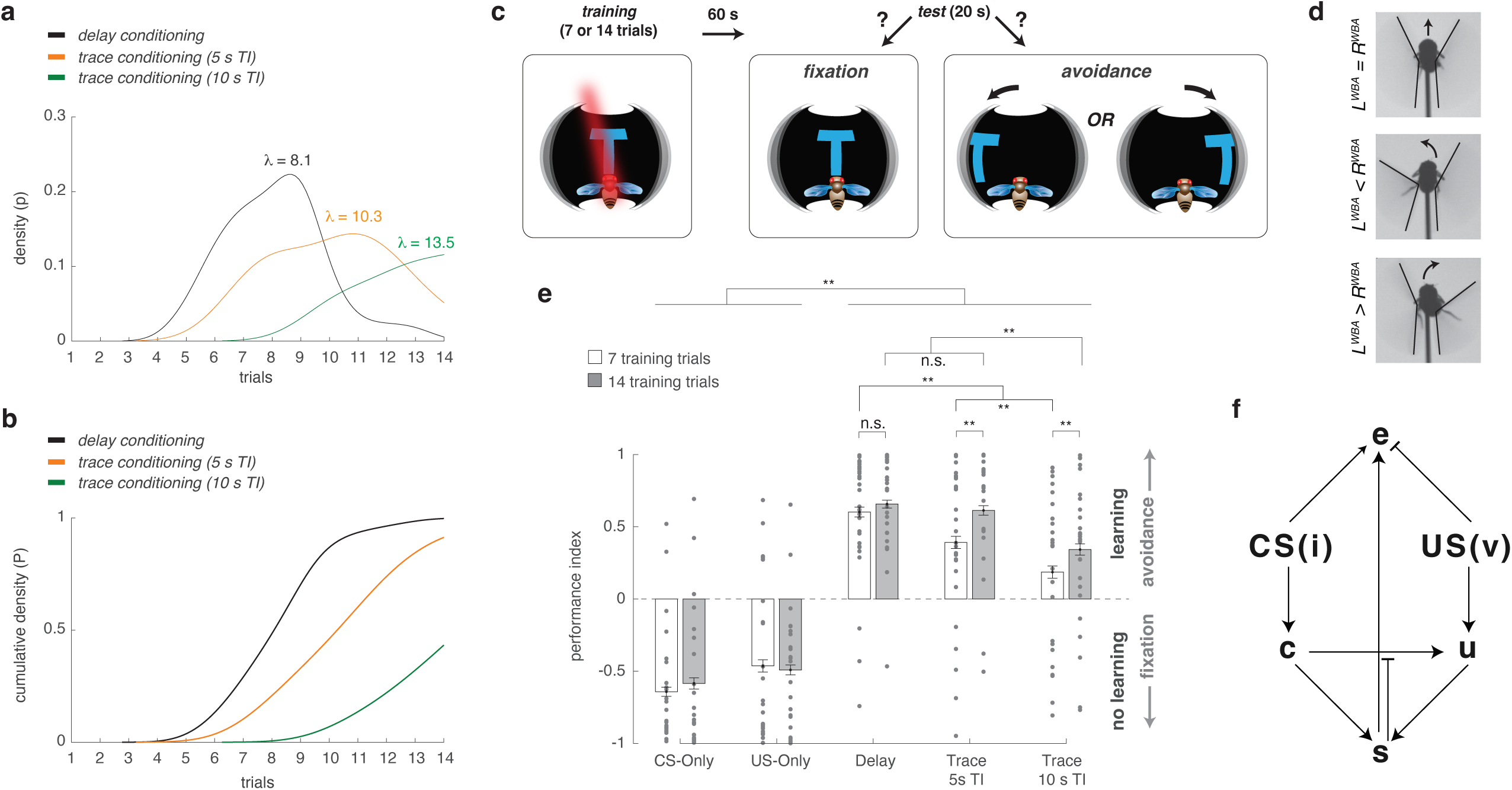
Tonic activity state shift predicts learning stability. **(a)** Change-point model applied to per-trial mean pre-CS baseline dF_ratio_/F_ratio_ activity (0-10 s) during delay conditioning (black, n = 42 flies), and trace conditioning with trace intervals of 5s (yellow, n = 11 flies), and 10 s (green, n = 15 flies). Shown are the probability distributions of each animal’s change point trial that partitions the data into two regions, minimizing the sum of the residual (squared) error of each region from its local mean. Maximum likelihood estimate of the α parameter of a Poisson distribution fit to each sample distribution. Change point modeling of this data is shown in **(Fig. 3i, S6e, S7e)**. **(b)** Cumulative distributions of each animal’s change point trial across learning conditions from **(a). (c)** Tethered-flight conditioning behavior assay illustrating upright-T (CS) paired with heat (US). Post-training (7 or 14 trials), fly’s closed-loop orientation towards (indication of no learning) or away (learning) from the CS is scored. **(d)** Wing-beat amplitude flying straight (top), yaw-turning counter-clockwise (center), and clockwise (bottom). **(e)** Conditioning performance indices (PI) for US-presentation only and CS-presentation only during training, compared to delay conditioning, and trace conditioning with 5 and 10 s trace intervals, for either 7 (white) or 14 (gray) training trials. (mean with s.e.m., n > 20 flies per group). Scatters represent single-fly PI. Groups compared using two-factor ART-ANOVA. n.s. indicates not significant, * is p-value < 0.05, ** p-value < 0.01. **(f)** Neuronal model integrating observed dopaminergic signals during delay and trace learning. Shown CS and US activating corresponding neurons *c* and *u*, which in turn jointly modify the shared synapse *s* such that the more their activity, the lower the weight of the synapse. These also activate an estimator neuron *e* that tracks, through its activity, the amount of time between CS-onset and US presentation. The higher the firing rate of *e*, the longer the interval. See Methods, Integrated neuronal model for further details.

We then measured behavioral performance in a subsequent test phase in which flies were presented with the conditioned visual cue in the absence of punishment. Consistent with prior studies of visual learning in *Drosophila*^27,39^, learned avoidance of the cue served as an index of associative memory formation (Fig. 5c–d). Behavioral acquisition closely paralleled the timing of the tonic state transition across conditions. Delay conditioning and 5 s trace conditioning both produced robust avoidance behavior by the end of training, corresponding to conditions in which most animals had undergone tonic state transitions. In contrast, 10 s trace conditioning resulted in delayed or incomplete tonic transitions accompanied by reduced avoidance of the CS within the same training period (Fig. 5e). Importantly, although the magnitude of avoidance was comparable between delay and 5 s trace conditioning, the timing of behavioral acquisition shifted in parallel with tonic transition timing. Across conditions, learned avoidance closely tracked the emergence of tonic state transitions.

Together, these results reveal a tight correspondence between tonic state transitions in PPM3-EB neurons and the acquisition of learned avoidance behavior, suggesting that this slower neural transition reflects a learned state linked to behavioral performance.

## Discussion

Our findings reveal that a single dopaminergic population in *Drosophila* exhibits multiple structured components of activity during associative learning across distinct timescales. PPM3-EB neurons displayed hallmark prediction-error responses, including a progressive shift in activity from punishment to the predictive visual cue, suppression following omission of expected punishment, and enhanced responses to stronger-than-expected punishment. At the same time, these neurons exhibited slower learning-state dynamics characterized by tonic baseline transitions and changes in perturbation variability that evolved across training, were jointly shaped by task structure, and tracked behavioral acquisition. Together, these findings suggest that dopaminergic activity in *Drosophila* integrates moment-to-moment prediction errors, temporal expectations, and slower learning-related dynamics within a single neural population.

Midbrain dopamine neurons in mammals are widely considered substrates of reward prediction-error signaling^3–6^, yet whether dopaminergic neurons simultaneously encode additional learning-related variables across longer timescales has remained unclear. Although theoretical and circuit models have proposed that *Drosophila* dopamine neurons could implement prediction-error computations^24–26^, direct physiological evidence has remained limited. Our findings extend this framework by showing that fly dopaminergic neurons exhibit several canonical features of prediction-error signaling, including transfer of responses from punishment to predictive cue, bidirectional responses to violated expectations, and sensitivity to expected punishment timing. These observations establish *Drosophila* as a tractable model for dissecting dopaminergic computations with trial-resolved and circuit-level precision.

Beyond event-locked prediction-error responses, we identified slower activity dynamics that evolved over training and followed temporal profiles distinct from phasic signaling. Previous studies have shown that dopamine prediction errors can be shaped by latent- or belief-state inference, in which reward expectations are computed over inferred environmental states^14,16,40^. In these frameworks, inferred state representations generally precede and influence prediction-error computation. By contrast, our data suggest that PPM3-EB activity itself contains separable components that evolve alongside learning and may reflect neural processes related to stabilization of predictive structure. Importantly, our findings do not demonstrate that PPM3-EB neurons explicitly encode latent task states. Rather, the abrupt transitions and resilience properties of tonic baseline activity are consistent with a learning-related reorganization of dopaminergic activity that emerges as cue–punishment contingencies become stabilized.

One possibility is that tonic baseline transitions reflect accumulation of evidence across trials, integrating prediction errors, temporal structure, and reinforcement history until a more stable activity regime emerges. Such transitions are unlikely to arise solely from prediction-error signaling or temporal relationships between cue and punishment. Learning-state inference is thought to integrate multiple sources of information^41,42^, raising the possibility that broader internal variables - including arousal, motivational state, attentional engagement, or locomotor state, contribute to when dopaminergic activity enters a stabilized regime. Future experiments combining circuit manipulation with behavioral-state measurements may help determine how these factors shape slow dopaminergic dynamics during learning.

Adaptive learning requires balancing stability and flexibility, maintaining reliable predictions when environmental contingencies are stable while remaining sensitive to change^43–45^. In this context, the dissociation between tonic baseline activity and perturbation variability may reflect complementary computational roles. Tonic baseline activity remained resilient to transient violations of learned contingencies, consistent with a more stable predictive regime, whereas perturbation variability rapidly emerged following initial punishment exposure and transiently increased when expectations were violated. These distinct resilience profiles raise the possibility that PPM3-EB dynamics contribute to balancing maintenance of learned associations with flexible updating when expectations fail.

Our results additionally suggest that dopaminergic activity carries information about the expected timing of punishment. Under trace conditioning, PPM3-EB activity progressively shifted toward the expected punishment time during the interval separating predictive cue and punishment, consistent with previous work implicating dopamine in temporal expectation and interval timing^37,38^. One possibility is that PPM3-EB neurons receive learning-dependent input from upstream temporal representations, analogous to hippocampal time cells^46^ or to the Mushroom Body Kenyon cell ensembles forming population clocks implicated in seconds-scale timing in *Drosophila*^47^. Alternatively, temporal estimation may emerge as an intrinsic component of dopaminergic learning itself, consistent with theories proposing that animals infer causal relationships from temporal structure^48^. Distinguishing between these possibilities will require future experiments probing how temporal information is represented and transformed across circuits during learning.

To synthesize these findings, we developed a minimal and interpretable circuit model capable of reproducing several key features of PPM3-EB activity (Fig. 5f; Methods, Integrated neuronal model)^3^. The goal of this model was not to provide a comprehensive mechanistic account, but rather to identify a minimal set of circuit interactions sufficient to generate the observed dynamics. In the model, dopaminergic activity, *u_T_(t)*, reflects the integrated output of sensory input from the predictive visual cue, *c_T_*, punishment input, *u_T_*, and a learning-dependent gating process governed by a plastic synaptic weight, *s_T_*,^49,50^ that evolves across trials *T*. As conditioning progresses, changes in *s_T_* alter how cue-related information contributes to dopaminergic activity, thereby linking learning history to trial-by-trial neural dynamics.

This simple architecture reproduced several core observations. First, the model captured the gradual reduction in punishment-evoked phasic responses as learning progressed. Early in training, high values of the learning synapse *s_T_* strongly weight punishment-driven activity, producing robust post-US responses in *u_T_(t)*. With learning, progressive depression of *s_T_* reduces punishment-dominated activity, resulting in the observed attenuation of punishment-evoked responses as expectations become established.

Second, the model reproduced the emergence of tonic baseline activity. During early training, elevated *s_T_* suppresses sustained dopaminergic activation, maintaining low tonic activity. As *s_T_* progressively decreases, cue-related input *c_T_(t)* increasingly contributes to *u_T_(t)*, eventually crossing a threshold that drives a transition toward elevated tonic baseline activity. This mechanism recapitulated the abrupt emergence of tonic state transitions observed experimentally.

Third, the model reproduced the progressive emergence of temporal expectation signals during trace conditioning through an interval estimator, *IT*, that updated trial-by-trial to improve predictions of the temporal delay between the predictive visual cue and punishment^51^. As learning progressed, the estimate of the cue–punishment interval became increasingly precise, allowing dopaminergic activity to progressively align with the expected punishment time. Importantly, because cue-related input *c_T_(t)* is weaker when punishment is temporally separated from the predictive cue, depression of *s_T_* proceeds more slowly, naturally reproducing the delayed acquisition observed during trace conditioning.

Finally, the model accounted for transient perturbations following violations of learned contingencies. Omission or enhancement of punishment on the final trial disrupted learned expectations, transiently increasing variability in *u_T_(t)* before the system returned toward its stabilized regime. Although intentionally simplified, this framework demonstrates how prediction-error responses, temporal expectation, and slower learning-state dynamics could emerge from interacting circuit processes operating across multiple timescales. As such, it provides a mechanistic starting point for future experiments aimed at identifying the cellular and circuit substrates underlying these computations.

In summary, our findings demonstrate that a single dopaminergic population in *Drosophila* simultaneously exhibits canonical features of prediction-error signaling, alongside slower learning-state dynamics, and temporal expectation signals during associative learning. These observations move beyond a view of dopamine as solely an event-locked teaching signal and instead suggest that dopaminergic activity integrates information across multiple timescales to shape learning under changing task demands. By establishing *Drosophila* as a tractable model for studying these interacting computations, this work provides a foundation for dissecting how prediction, temporal structure, and slower learning dynamics are jointly implemented in neural circuits.

## Methods

### Fly stocks

*Drosophila melanogaster* flies were raised and maintained on a standard cornmeal-agar medium at 25°C, 60% humidity, on a 12-h light-12-h dark cycle. All experimental flies were non-mated females 3-10 days old, isolated post-eclosion at 25°C, and tested 0-3 h before the onset of the dark cycle. *c346-Gal4* (30831), *20XUAS-GCaMP7f* (79031) and *10XUAS-myr-tdTomato* (32222) were all obtained from the Bloomington Drosophila Stock Center.

### Virtual reality tethered-flight behavior assay

Female flies were cold-anesthetized (4°C) and tethered to a stainless steel minutien pin (0.2 mm rod diameter, Fine Science Tools) using UV-cured glue, curing for at least 30 seconds. After a recovery period of at least one hour, these tethered flies were suspended within a high-speed projector-based spherical virtual-reality flight arena. The exterior of the 2 in diameter sphere was coated using a rear-projection medium (Screen Goo, Goo Systems Global) allowing the presentation of dynamic computer-generated stimuli and create an immersive virtual environment.

The tethered-flight arena followed the basic protocol outlined previously to generate an immersive visual surround^27^. Briefly, a high-speed projector (Texas Instruments, LightCrafter 4500 EVM) projected three virtual views of a scene onto the sphere, the first head on, the other two lateral views reflected off two 2-inch mirrors (Edmund Optics) that were placed on either side to the sphere. This produced a virtual-reality display encompassing around 330° of azimuth and around 85° of elevation. Custom software written in Microsoft Visual C++ using a 3D graphics programming library OpenSceneGraph allowed us to warp and display images on a curved spherical projection screen. The projector was set to display 6-bit images at a resolution of 912 x 1140 at 300 Hz and emitted a blue light which was filtered through a 450-nm long-pass emission filter (62-982, Edmund Optics). Calibration of light intensity was performed using a SpectraScan PR-701S spectroradiometer and the intensity was controlled by altering the current of the projector LED, ultimately set to 10% of the maximum power. The overall radiance inside the spherical area for an all-ON stimulus was set to be 0.4 W m^-2^ sr^-1^, to match the light intensity during natural crepuscular sunset.

To illuminate the fly and enable measurement of its wingbeat amplitude, a custom infra-red diffused backlight of 49 LEDs (Vishay, 2 in x 2 in, 880 nm) was positioned below the sphere. To capture the wingbeats, a high-speed camera (FL3-U3-13Y3M-C, Point Grey Research) with attached lens (Tamron macro lens, f/2, 60 mm) and an infrared pass-only filter (850 nm, Edmund Optics) was positioned above the sphere. The amplitudes of the fly’s left- and right-wingbeats were computed in real-time using custom software written in Microsoft Visual C++ using the machine vision OpenCV library at 200 Hz, allowing us to present visual stimuli both with and without yaw-turning closed-loop feedback.

Using a focusable dot infra-red laser (808 nm, maximum power 350 mW, Roithner LaserTechnik), we were able to deliver a heat punishment which was pre-calibrated to raise the fly’s body temperature from ambient room temperature (25 °C) to either 35 °C or 40 °C within 0.5s. Laser power to achieve these temperature increases were determined by inserting the top of a hypodermic needle thermocouple probe (HYP1-30-1/2-T-G-SMPW-M, Omega). An 850 nm long-pass dichroic beamsplitter (Edmund Optics) was placed in between the display sphere and the high-speed camera above it to direct the punishment laser light beam down on to the fly while still allowing for the camera above to detect the wing-beat image.

### Conditioning protocols and learning performance

During the training phase, we presented an upright-T shaped bright image over a dark background to naïve flies (not exposed to any conditioning previously). The T-shaped visual cue was displayed in the frontal visual field of the fly and was held fixed during presentation. The T-shape measured 40° vertically and horizontally, with the bar of the T being 14° wide. Delay and trace conditioning (5 and 10 s trace intervals) involved presenting the CS and US for either 7 or 14 training trials, with each training trial lasting 60 s with a 30 s inter-trial-interval. US-only and CS-only presentations (for the same number of training trials) served as learning controls. Post-training, a single test of learning was performed for each conditioned fly. A 60 s gap was introduced between the end of the last training trial and start of the test sequence.

During the test phase, the CS was presented in the fly’s frontal visual field in closed-loop mode for 20 s. Learning performance was quantified using a performance index (PI), calculated based on the fly’s orientation relative to the CS. Specifically, we measured the time the fly spent fixating on the CS in the center (frontal) quadrant (*t_a_*) and the time spent orienting away from the CS while holding it laterally in the left or right quadrants (*t_b_*). The PI was calculated as *(t_b_ - t_a_)/(t_b_ + t_a_)*. A positive PI (PI > 0) indicated successful learning, reflecting longer fixation away from the CS, whereas a negative PI (PI < 0) indicated greater fixation toward the CS. A PI of 0 indicated no preference, corresponding to equal time spent fixating toward and away from the CS. To ensure reliable behavior measurements, only experiments in which flies spent at least 50% of the test period (>10 of 20 s) fixating within the defined quadrants were included in the analysis.

### Fly brain window surgery

A custom flyholder was used to immobilize the fly head under an imaging device and expose its brain for optical recordings (^52^). The flyholder consisted of two parts – a 3D printed plastic frame, and a soft annealed stainless-steel shim (with a fly brain sized hole) folded and glued with epoxy to fit the contour of the frame. After gluing the metal shim to the holder, charcoal primer and paint were applied to the bottom side of the shim to minimize reflections during wing-beat tracking.

Our fly mounting and surgery procedure is as described - cold-anesthetized flies (4 °C) were positioned ventral side up in a fly-sized divot machined in a custom-made brass block. The first pair of legs (T1) were cut at the first segment, and the middle and rear pairs (T2 and T3) were removed completely. The proboscis was gently pushed into the head capsule, and a small drop of UV-cured glue was applied to fix it in place. The fly was then flipped over, dorsal side up, and a small drop of UV-cured glue was applied in the gap between the head and thorax, thereby tilting the fly head slightly upwards. The flyholder was then positioned and glued to the head, following which, the holder and the fly were removed from cold-anesthesia and allowed to recover.

After recovery (determined by the fly exhibiting flight), the flyholder was filled with saline to fully cover the head. 1x saline containing 103 mM NaCl, 5 mM TES, 8 mM Trehalose, 10 mM Glucose, 26 mM NaHCO_3_, 1 mM NaH_2_PO_4_, 1.5 mM MgCl_2_, and 3mM CaCl_2_, with a pH = 7.3 was prepared and used. With the head covered in saline, the cuticle, air sacs and fat bodies were removed by hand dissection using fine forceps (Dumont 5SF, Fine Science Tools, or tip-size A, re-sharpened at Corte Instruments). An incision was made first along the posterior side of the head, then along both lateral edges, and finally along the anterior side to remove the cuticle.

### Two-photon *in-vivo* fluorescence imaging

The tethered flight behavior assay described above was coupled with a two-photon microscope with some modifications. First, a cylindrical acrylic display (4 in diameter, 3.5 in height, ∼330° of azimuth and ∼85° of elevation) with an adhesive rear-projection film was used for visual stimulus presentation. The display was tilted downwards (pitch angle of ∼20°) relative to the flyholder positioned in it. The projector used for visual stimulus presentation was identical to the tethered-flight assay except for a 447/60 bandpass filter (Chroma) that was placed in the light path to avoid any bleed through of visual stimulus light into the imaging recordings.

A collimated fiber-coupled 850 nm IR LED (M850F2, M28L01, F240SMA-850, Thorlabs), placed directly under the flyholder and behind the fly, was used to illuminate it for accurate positioning, the image of which was captured using a high-speed IR sensitive camera (FL3-U3-13Y3M-C, Point Grey Research). The IR laser punishment delivery apparatus (808 nm, maximum power 350 mW, Roithner LaserTechnik) was identical to the tethered flight assay and delivered from underneath the fly.

Our imaging experiments were performed using a Bruker Ultima Investigator multiphoton microscope with a Nikon 40x NIR Apo objective water-immersion lens (0.8 N.A., pixel size 0.27 x 0.27 μm, FOV 69.1 x 69.1 μm). GaAsP photomultiplier tubes (H10770, Hamamatsu) band-passed with either et525/70m-2p or et595/50m-2p emission filters (Chroma) and a t565lpxr dichroic beam-splitter (Chroma) were used for simultaneous acquisition. A 920 nm mode-locked Ti:Sapphire laser (Mai Tai, Spectra Physics) provided a maximum power of 15 mW on the sample. Resonant scanning galvos and a high-speed Z-piezo (Bruker, max range, 400 μm) were used for imaging a 6-plane volume of the EB at a rate of ∼6 Hz (image resolution of 256×256 px, 5 μm spacing between imaging planes).

Multi-plane 3D volumes of calcium-dependent GCaMP and calcium-independent tdTomato fluorescence from the same neural populations were simultaneously captured with dual GaAsP PMTs.

### Calcium fluorescence quantification

We first generated a maximum intensity projection (MIP) sequence for each pixel (256×256 px) in the 3D volume for GCaMP and tdTomato channels. Each MIP sequence was filtered using a 3×3 median spatial filter followed by 2D gaussian smoothing with a square gaussian kernel standard deviation of 1, to smooth acquisition noise. Next, the tdTomato channel sequence was then spatially aligned using a fast-normalized cross-correlation template matching algorithm based on the first MIP frame. The same transformations were then applied to the MIP sequence of the GCaMP channel to ensure pixel-pixel correspondence between frames of the two sequences.

We then created a binary mask of labeled neurons, segmenting foreground (neural) pixels from background. The mask was created by applying adaptive thresholding to each frame of the tdTomato channel. The calcium transient was then measured as a ratio, of average green calcium-dependent fluorescence (GCaMP) over red calcium-independent fluorescence (tdTomato) within the masked region.

Change in fluorescence intensity levels were then determined by first applying a Savitzky-Golay filtering (order of 3, frame length of 7) to the raw intensity time-series data, then computing the difference of maximum F values from a baseline level: *ΔF/F = (F - F_baseline)/F_baseline*. Baseline levels were computed by averaging the lowest 25 % intensity frames during the baseline period (5 s) of the first trial before CS presentation.

### Peak detection analysis

Local maxima peaks, defined as points where the value is higher than all nearby points, indicating a rise-then-fall pattern, were determined for fluorescence recordings. Minimum peak height and prominence thresholds of 5-standard-deviations of the corresponding baseline *ΔF/F* level were used to detect significant peaks in fluorescence. Peak prominence measured how much a peak stood out from the surrounding signal, defined as the vertical distance from the peak to the lowest contour line that encircled it and no higher peaks.

### Change-point analysis

To identify abrupt transitions in dopamine activity during learning, we performed change-point analysis. Trial-by-trial tonic baseline activity and perturbation-related variability were analyzed separately for each animal. The algorithm identifies points in a time series where statistical properties change by minimizing within-segment squared error under a piecewise-constant model. We used a single change-point model to test the hypothesis that learning-related activity undergoes one dominant transition between distinct states. The location of the change point was estimated independently for each animal and defined as the trial at which the model detected the largest shift in mean activity or variability. To assess consistency across animals, estimated change points were compared across the population and visualized relative to learning progression.

### Hidden state modeling of dopamine activity during learning

Trial-wise dopamine activity was quantified from calcium imaging recordings by computing the mean fluorescence signal (*ΔF/F*) within predefined analysis windows for each conditioning trial. To account for differences in overall signal amplitude across animals, activity values were normalized within an animal by z-scoring across learning trials.

To model latent transitions in dopamine activity during learning, we used a two-state Hidden Markov-switching model. The model consisted of two latent states corresponding to pre-learning and learned regimes. State transitions were initialized using a left-right (“sticky”) transition structure designed to favor persistent occupancy of the current state and minimal backward transitions. The a priori transition probability matrix was initialized as:

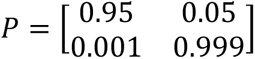

where rows represent the current state and columns represent the subsequent state. This initialization reflects the assumption that animals begin in a pre-learning state, transition infrequently into the learned state, and rarely revert to the pre-learning state once learning has occurred. Model parameters, including state-transition probabilities and state-specific emission parameters, were subsequently estimated from the data by maximum likelihood.

State-specific emissions were modeled as Gaussian processes with independent means and variances. For each trial, smoothed posterior state probabilities were computed from the fitted model. The probability of occupying the learned state was used to quantify the emergence of learning over trials. For each animal, the learning transition trial was defined as the first trial at which the posterior probability of the learned state exceeded 0.8 for at least two consecutive trials.

To assess the effects of prediction error, activity on the probe trial was compared across conditions (normal conditioned stimulus–unconditioned stimulus pairing, omission of the unconditioned stimulus, and increased unconditioned stimulus magnitude). Probe-trial activity was evaluated relative to the learned-state emission distribution by computing a normalized deviation:

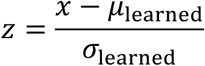

where *x* is the probe-trial activity and *μ*_learned_ and *σ*_learned_are the estimated mean and standard deviation of the learned-state emission. Log-likelihood ratios comparing the learned and pre-learning state emissions were additionally computed for each probe trial. Condition effects were tested using linear models, followed by analysis of variance. Trial-to-trial changes in tonic activity were quantified as the difference between probe-trial activity and the final learning trial.

### Analysis of perturbation variability

To quantify perturbation variability in dopamine activity following punishment delivery, we computed the standard deviation (SD) of *ΔF/F* during the post-outcome period for each trial. Because SD values were positively skewed, values were log-transformed prior to analysis and subsequently normalized within animal by z-scoring across learning trials.

Latent-state analyses of perturbation variability were performed using the same two-state Hidden Markov-switching framework described above. Probe-trial variability was evaluated relative to the learned-state emission distribution using normalized deviations and log-likelihood ratios. To quantify probe-induced changes in perturbation variability, trial-to-trial differences were computed relative to the preceding learning trial.

### Comparison of tonic and variability signals

To compare the effects of prediction errors on tonic baseline activity and perturbation variability directly, normalized deviations from the learned-state distribution were analyzed jointly using a linear model containing signal type (mean baseline activity vs. standard deviation) and probe condition as factors. Interaction terms were used to test whether prediction error manipulations differentially affected tonic and variability components of dopamine signaling.

### Correlation between learning and variability modulation

To assess the relationship between learning and prediction error-induced variability changes, learning metrics derived from the latent-state model were correlated with probe-induced changes in spontaneous variability across animals. Learning strength was quantified as the posterior probability of occupying the learned state on the final learning trial. Correlations were computed using Spearman rank correlation coefficients.

### Statistics and reproducibility

We performed all post-acquisition analyses using Matlab R2024a. Summarized data were represented either as means (with s.e.m) or boxplots, with center (median), and edges (inter-quartile range).

When comparing more than two groups, Kruskall-Wallis and post-hoc unpaired two-sided Mann–Whitney U tests with bonferroni corrected multiple comparisons were used. Two group comparisons were performed using unpaired two-sided Mann Whitney U tests. For testing equality of variance between groups with similar means, the Brown-Forsythe test was used. Variance specific Holm correction was used for multiplicity correction.

Behavioral and neurophysiological data with a two-factor structure (across trials and conditioning variants) were analyzed using two-factor aligned-rank-transform analysis of variance (ART-ANOVA). Single-factor comparisons (across trials for same conditioning paradigm) were performed using Friedman’s repeated measures ANOVA. Sample size estimates based on medium to large effect sizes (Cohen’s d), were calculated a priori, during the experimental design stage. Final sample sizes ensured a test power of at least 0.8. Experiments were performed non-blinded. Single fly metrics are overlaid in all figures as scatters.

### Integrated neuronal model

Here, we present a simplified rate coding model of learning and prediction error signaling based on earlier studies^3,21^. The model captures the elements of the learning process as shown in **Fig. 5f**. There are 2 input variables - *i*, the T-shaped predictive visual cue and *v,* the valence (punishment in this case). These, in turn, activate neurons *c* and *u* in the brain that channel the activity of *i_t_* and *v_t_* onto learning processes. *u* represents the activity of the dopaminergic neuron, and *c* represents the CS. *c* and *u*, in turn, coincidentally activate the learning synapse *s*. Here, *s* is synaptic weight and takes values between 0 and 1. The co-incident detection of CS and US depresses the synaptic weights^50,53^. In the model, *u* captures the following characteristic of the learning process - it learns to estimate the time interval between CS and US, increases tonic activity levels with progression in learning, and captures the decrease and increase in noise as learning improves or worsens with each trial.

The model also includes two intermediate variables that do interval estimation^51^. First, *e* is an estimator neuron that through its activity represents the elapsed time from the start of the CS and US presentation. The higher the value of *e*, the longer the time interval. Second, *IT* is the interval time estimate which is the result of differencing the interval for the current trial *T*, and the previous (*T-1*) trial’s interval. Both these variables approach an asymptotic value as learning progresses.

There are two temporal variables, *t*, depicting time within an interval and *T*, the trial number. For brevity, all variables are written with a subscript, *t* or *T*, indicating timescale on which they evolve. For example, *e_t_* denotes estimator neuron activity across time *t* within a trial. It could also be written as *e_T_(t)*. The same goes for *e_T_*, which is the estimator neuron’s activity at the end of trial *T* or *e_T_ = max[e_T_(t)]*. The same goes for variables *c, u, s*, *i,* and *v*. Finally, *k* and λ control the rates of activation or inhibition, and the rates of decay of the variables.

The equations that capture the workings of the model are as follows.

*e_t_*, the time estimator,

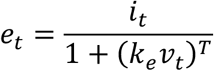

*IT*, the interval time,

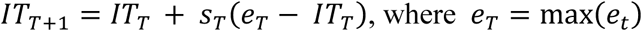

*s_T_*, the synapse that captures the CS-US association,

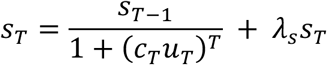

When there is coincident activity, the synapse strength goes down. Otherwise, the strength either stays the same or increases by a small amount.

*c_t_*, the neuron that reflects the CS,

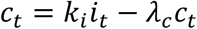

*u_t_*, the dopaminergic neuron activity,

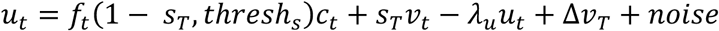

where *thresh_s_* is a scalar value for activation and deactivation of a synapse *s*, *f_t_* is the general threshold function, 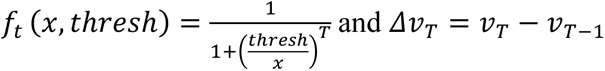

The first term, *f_t_* (1 – *s_T_*, *thresh_s_*)*c_t_* captures the threshold activation for synapse *s* that leads to an increase in tonic activity. The second term *v_t_s_T_* captures the bump in activity when a punishment is presented in early trials. With learning *s_T_* decreases and this has a lower effect on *u_t_*. The third term *λ_u_u_t_* captures the degradation of *u_t_*. In this rate coding formulation, *α* is very low and is not critical to the model but included for completeness. *Δv_t_* captures the change in activity due to the sign of the valence. *noise* comprises two components: *interval estimator* and *post-US-noise*. Estimator noise precedes US onset and reflects variability in the learned estimate of the CS-US interval. During early trials, the dopaminergic neuron underestimates the CS-US interval, resulting in repeated activity peaks before US onset. As learning progresses, the interval estimate gradually converges toward the true interval.

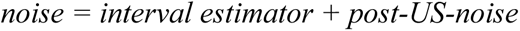

*interval estimation:* 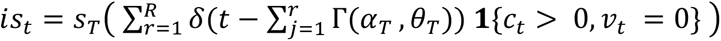

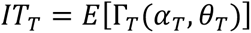

*cv_T_* = *cv_T_*_-1_*s_T_, where cv*_0_ = 0.1, and *cv* is the coefficient of variation

*post-US-noise: noise_p_* = *f_t_* (*s_T_*, *thresh_s_*) **1** {*i_t_* = 0, *v_t_* = 0)

An explanation for interval estimation time (*IT*) is warranted. The estimate of the CS-US interval is updated after each trial based on the discrepancy between the current interval estimate and the observed interval, together with information carried forward from previous trials. Within a trial, presentation of the CS drives activity in the estimator, *e_t_*, which accumulates while the CS is present. This accumulated activity is used to update the estimate of the interval separating the CS and US onset. Early in training, the estimated interval is imprecise and typically shorter than the true interval. As learning progresses and *s_T_* evolves, the interval estimate becomes increasingly accurate and converges toward the CS-US interval.

The learned interval estimate is then used to generate anticipatory dopaminergic activity centered on the predicted time of US occurrence. Specifically, anticipatory activity is modeled using a Gamma distribution whose expected value is given by *IT_T_*. The distribution is parameterized by shape and scale parameters (*α_T_*, *θ_T_*). Following the first trial, the initial estimates *IT_T_* and *cv_1_* are obtained, allowing the distribution parameters to be calculated. As learning proceeds, the variance of the estimate decreases, producing increasingly precise timing of anticipatory dopaminergic activity.

The equations presented here are intended as a simplified rate-coding description of the proposed mechanism rather than a detailed biophysical model. Several variables, including the elapsed-time estimator *e_t_*, are represented using direct rate-based equations that do not explicitly incorporate memory. More biologically realistic implementations could employ recursive update rules and leaky accumulator architectures that generate similar functional behavior. Although such alternatives are not mathematically equivalent to the equations used here, they serve analogous computational roles and are expected to produce qualitatively similar model behavior. For simplicity and analytical clarity, we therefore use the reduced rate-coding equations shown in the equations above.

## Data availability

Datasets generated as part of this study will be made available from the corresponding author on request.

## Code availability

All source codes for the different assays used and analysis routines will be made available for download from the public repository at https://github.com/dgrover/flyRPE.

### Acknowledgements

We thank R Greenspan, K Asahina and members of the Grover laboratory for discussions and comments on the study. This work was supported by Air Force Office of Scientific Research grants FA9550-23-1-0024 and FA9550-25-1-0299 to DG.

## Author contributions

WC performed the experiments. WC and DG designed the study and performed data analysis. SS developed the neuronal model. DG supervised the study. WC, SS and DG wrote the manuscript. All authors reviewed and approved the final manuscript.

## Supplementary figure

**Figure S1.**
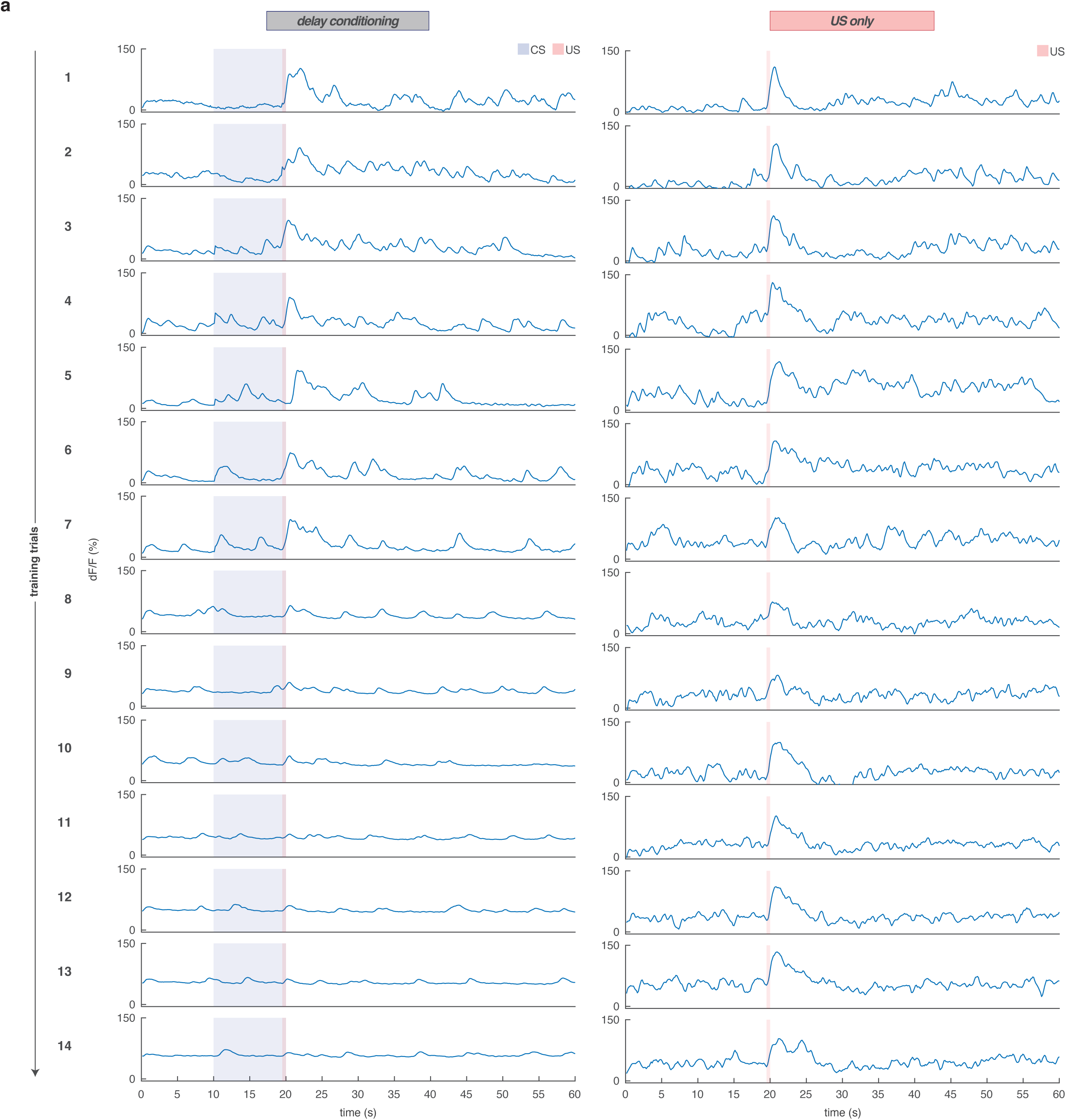
Ratiometric calcium imaging of EB-projecting PPM3 dopaminergic neurons during US-Only condition do not show phasic prediction error-like activity and tonic activity changes. **(a)** Ratiometric calcium imaging of a *c346-Gal4>>UAS-GCaMP7f;UAS-myr-tdTomato* female during delay conditioning (left) and US-only (right). Shown, dF_ratio_/F_ratio_ activity (trials 1 thru 14).

**Figure S2.**
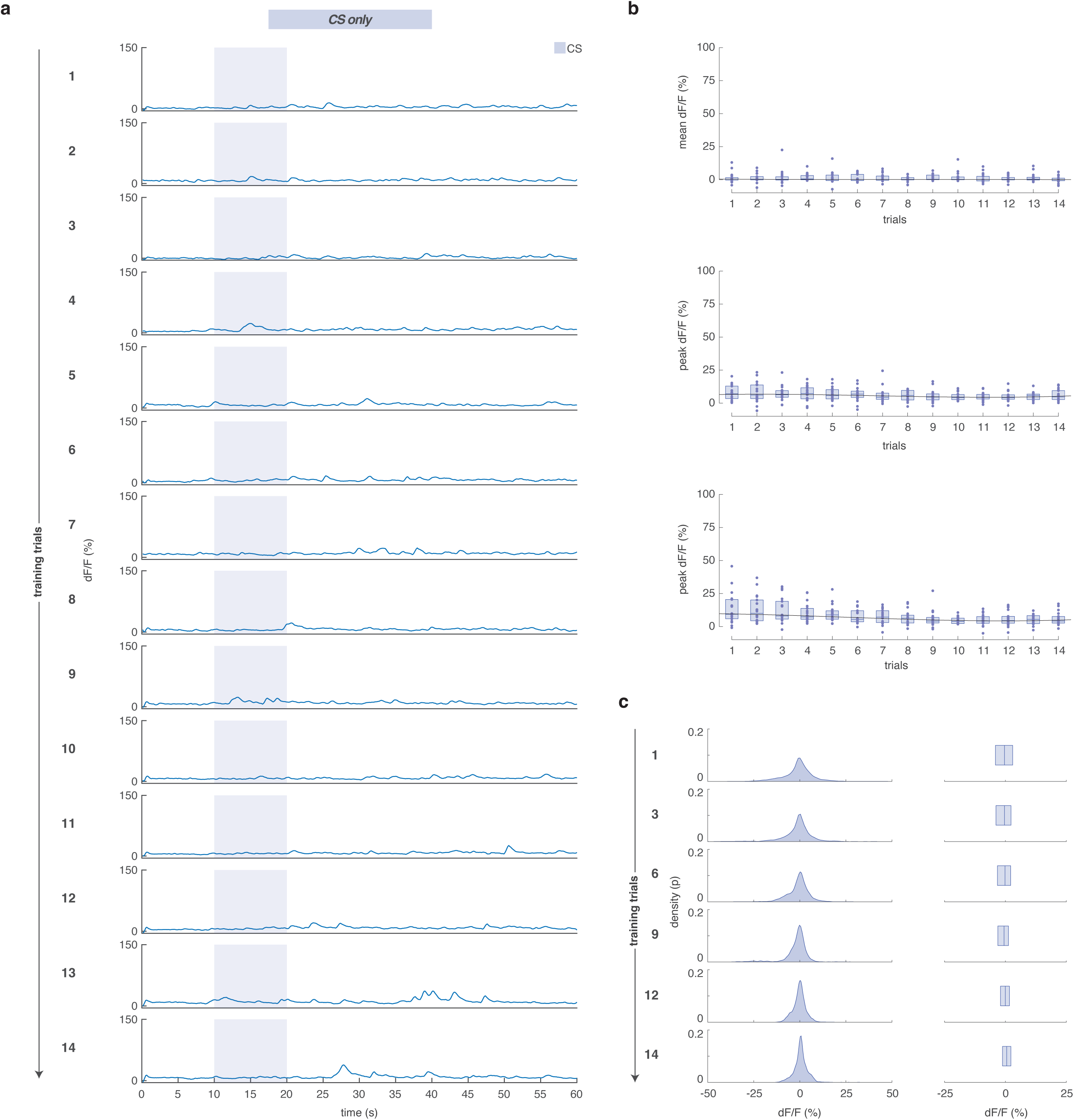
Ratiometric calcium imaging of EB-projecting PPM3 dopaminergic neurons during CS-Only condition do not show phasic prediction error-like activity and tonic activity changes. **(a)** Ratiometric calcium imaging of a *c346-Gal4>>UAS-GCaMP7f;UAS-myr-tdTomato* female during CS-only condition. Shown, dF_ratio_/F_ratio_ activity (trials 1 thru 14). **(b)** Top row, mean dF_ratio_/F_ratio_ activity during the pre-CS baseline period (0-10 s) for CS-only condition (n = 18 flies). Middle and bottom rows are peak dF_ratio_/F_ratio_ activity during CS (10-20 s), and post-US (20-25 s). Shown are quadratic (second degree) polynomial curve fits through median dF_ratio_/F_ratio_ activity. **(c)** Distributions of perturbation variability (relative to pre-CS baseline mean) from 25-60 s (5 s after delay conditioning matched-US timing). Shown are kernel probability density estimate (left) and boxplot (right) representations for trials 1, 3, 6, 9, 12, 14. Boxplot center (median), and edges (IQR). Scatters represent single-fly metrics.

**Figure S3.**
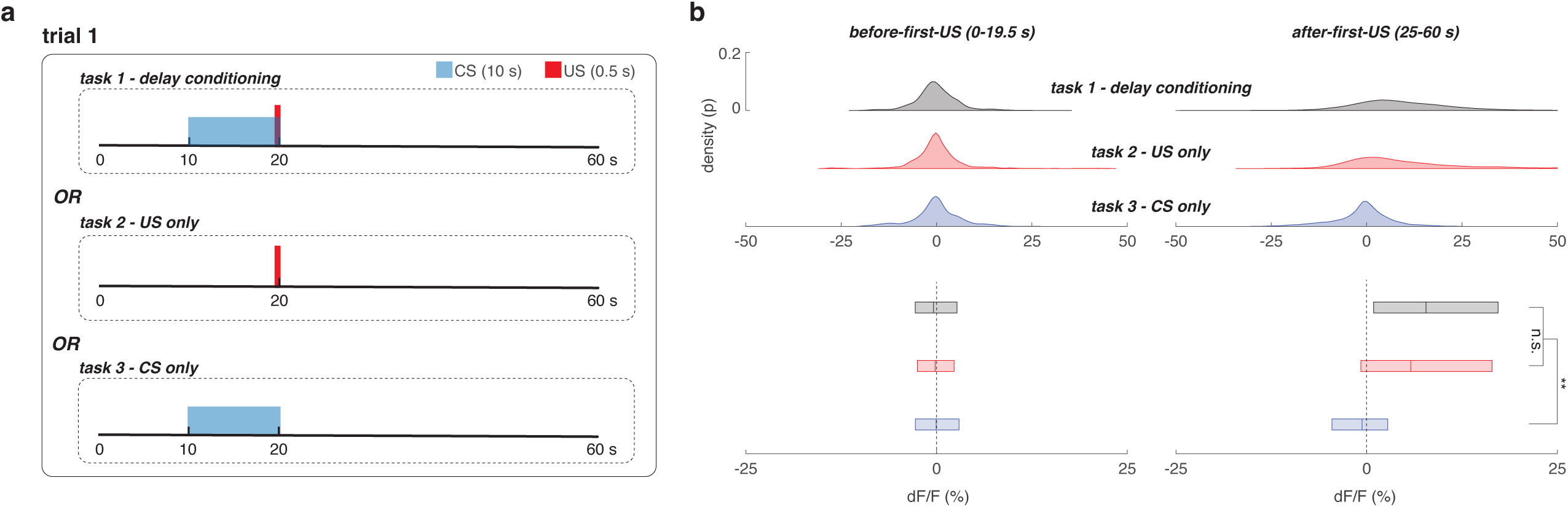
Initial punishment gates the perturbation variability of EB-PPM3 neurons regardless of learning. **(a)** Illustration of three conditioning tasks tested - delay conditioning wherein the predictive visual cue was paired with the punishment (top), US-only where only the punishment was given (middle), and CS-only where only the visual cue was presented without the punishment (bottom). **(b)** Comparison of the distributions of perturbation variability (relative to pre-CS baseline mean of trial 1) before presentation of first US (left column, 0-19.5 s) in trial 1 as shown in (a) and 5 s after US (right column, 25-60 s). Shown are data from trial 1 of delay conditioning (black, n = 42 flies), US-only (red, 41 flies), and CS-only (blue, n = 18 flies). Shown are kernel probability density estimate (top row) and boxplot (bottom row) representations. Equality of group variances was tested using Brown-Forsythe variant ANOVA with Holm multiplicity correction. Boxplot center (median), and edges (IQR). n.s. indicates not significant, * is p-value < 0.05, ** p-value < 0.01.

**Figure S4.**
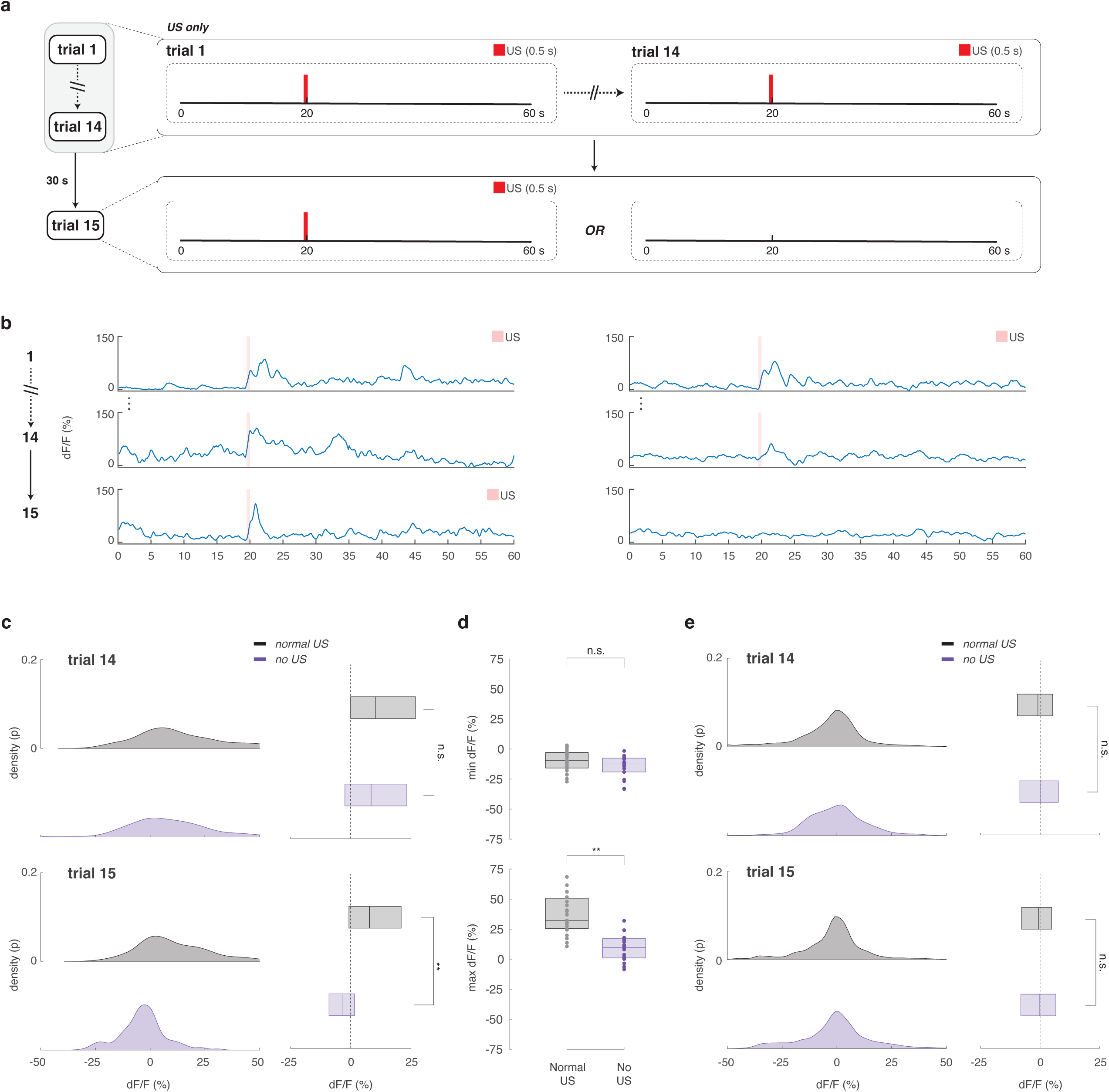
Ratiometric calcium imaging of EB-projecting PPM3 dopaminergic neurons during US-Only condition does not show the dopamine dip in response to the omission of punishment. **(a)** Illustration of the experimental design used as a control test for dopamine prediction error signaling - 14 trials of US-only task, followed by a probe trial 15 that tests one of two conditions – typical US-only trial (left), or omission of US. **(b)** Ratiometric calcium imaging of a *c346-Gal4>>UAS-GCaMP7f;UAS-myr-tdTomato* females for the two conditions described in **(a)**, during typical US-only (left), and 14 trials of US-only followed by omission of US in trial 15 (right). Shown, dF_ratio_/F_ratio_ activity (trials 1, 14, and probe trial 15). **(c)** Comparison of the distributions of activity (relative to matched-pre-CS 0-10 s baseline mean) during the post-US interval (20-25 s) for trial 14 (top row) and probe trial 15 (bottom row) for the two conditions in **(a)**. Shown are kernel probability density estimate (left column) and boxplot (right column) representations. **(d)** Minimum (top) and maximum (bottom) dF_ratio_/F_ratio_ activity (relative to matched-pre-CS 0-10 s baseline mean) during post-US interval (20-25 s) in probe trial 15 for **(b)**. **(e)** Comparison of the distributions of perturbation variability (relative to matched-pre-CS 0-10 s baseline mean) 5 s after US (25-60 s) for trial 14 (top row) and probe trial 15 (bottom row) for the two conditions in **(a)**. Shown are kernel probability density estimate (left column) and boxplot (right column) representations. Equality of group variances was tested using Brown-Forsythe variant ANOVA. In **(c-e)**, shown are data from US-only (black, n = 23 flies), and omission of US condition (purple, 18 flies). Boxplot center (median), and edges (IQR). Scatters represent single-fly metrics. Group distributions compared using unpaired two-sided Mann–Whitney U tests. n.s. indicates not significant, * is p-value < 0.05, ** p-value < 0.01.

**Figure S5.**
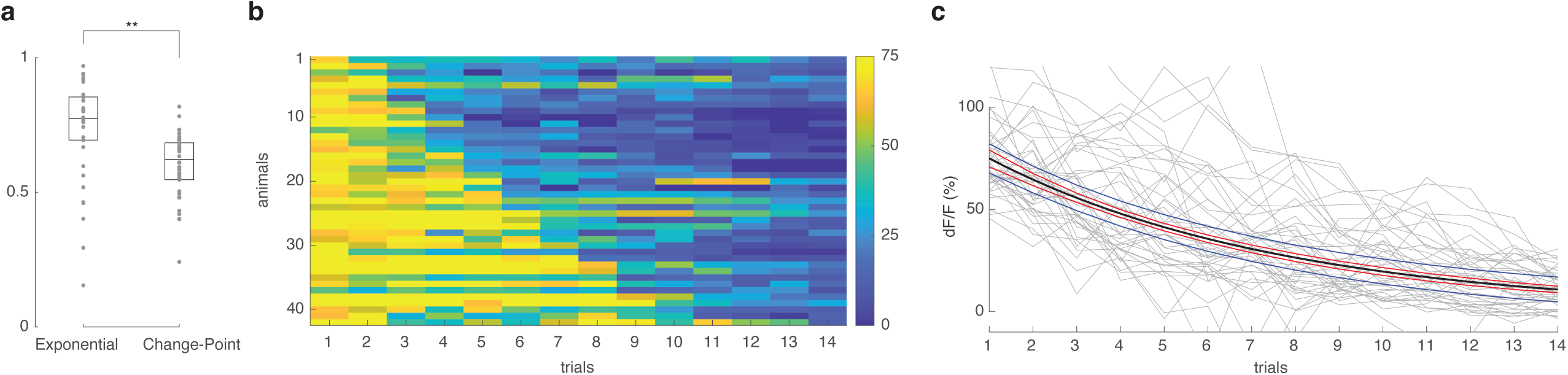
EB-PPM3 Post-US PE calcium activity exhibits an exponential decay. **(a)** R-squared goodness of fit of exponential and change-point models applied to per-trial peak dF_ratio_/F_ratio_ post-US activity (20-25 s) during delay conditioning (n = 42 flies). **(b)** Heatmap of modeled data from **(a)** in ascending order of each animal’s change point trial that partitions the data into two regions, minimizing the sum of the residual (squared) error of each region from its local mean. **(c)** Gray lines are the same data as **(b)**. Single-term exponential curve-fit (black), with 95% upper- and lower-simultaneous observation (blue) and functional (red) prediction bounds. Boxplot center (median), and edges (IQR). Scatters represent single-fly metrics. Groups compared using unpaired two-sided Mann–Whitney U tests. n.s. indicates not significant, * is p-value < 0.05, ** p-value < 0.01.

**Figure S6.**
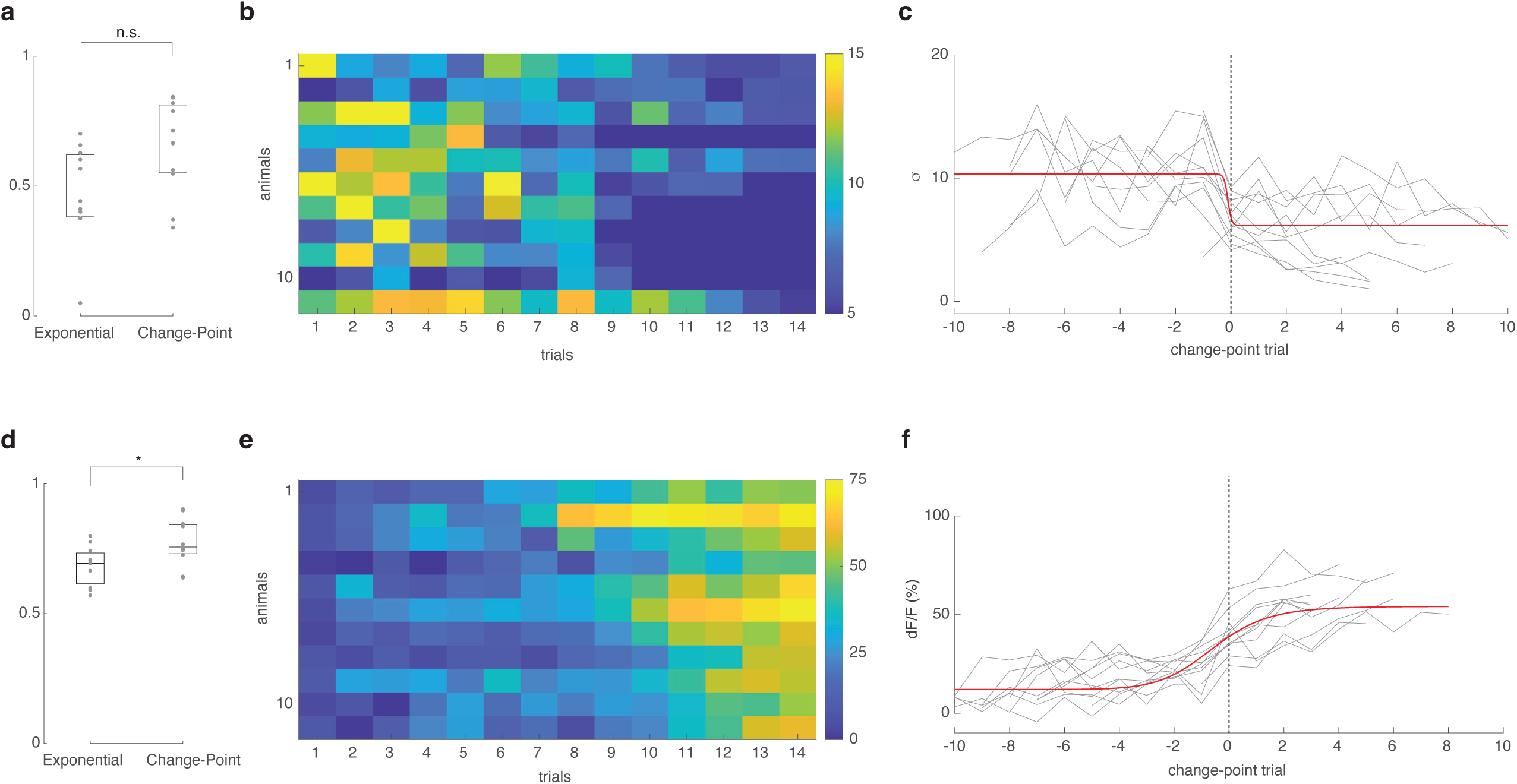
Ratiometric calcium imaging of EB-projecting PPM3 dopaminergic neurons during trace conditioning reveals that a 5s trace interval between CS and US delays state transition trials. **(a)** R-squared goodness of fit of exponential and change-point models applied to per-trial standard deviations of dF_ratio_/F_ratio_ perturbation variability (relative to pre-CS baseline mean) 5 s after US (30-60 s) during trace conditioning with a 5 s trace interval (n = 11 flies). **(b)** Heatmap of modeled data from **(a)** in ascending order of each animal’s change point trial that partitions the data into two regions, minimizing the sum of the residual (squared) error of each region from its local mean. **(c)** Same data as **(b)**, trial aligned to center (trial 0) on each animal’s respective change point trial (gray lines). Red curve indicates sigmoidal curve fit to the per-trial median of the change point trial-aligned data. **(d)** R-squared goodness of fit of exponential and change-point models applied to per-trial mean pre-CS baseline dF_ratio_/F_ratio_ activity (0-10 s) during trace conditioning with a 5 s trace interval (n = 11 flies). **(e)** Heatmap of modeled data from **(d)** in ascending order of each animal’s change point trial that partitions the data into two regions, minimizing the sum of the residual (squared) error of each region from its local mean. **(f)** Same data as **(e)**, trial aligned to center (trial 0) on each animal’s respective change point trial (gray lines). Red curve indicates sigmoidal curve fit to the per-trial median of the change point trial-aligned data. Boxplot center (median), and edges (IQR). Scatters represent single-fly metrics. Groups compared using unpaired two-sided Mann–Whitney U tests. n.s. indicates not significant, * is p-value < 0.05, ** p-value < 0.01.

**Figure S7.**
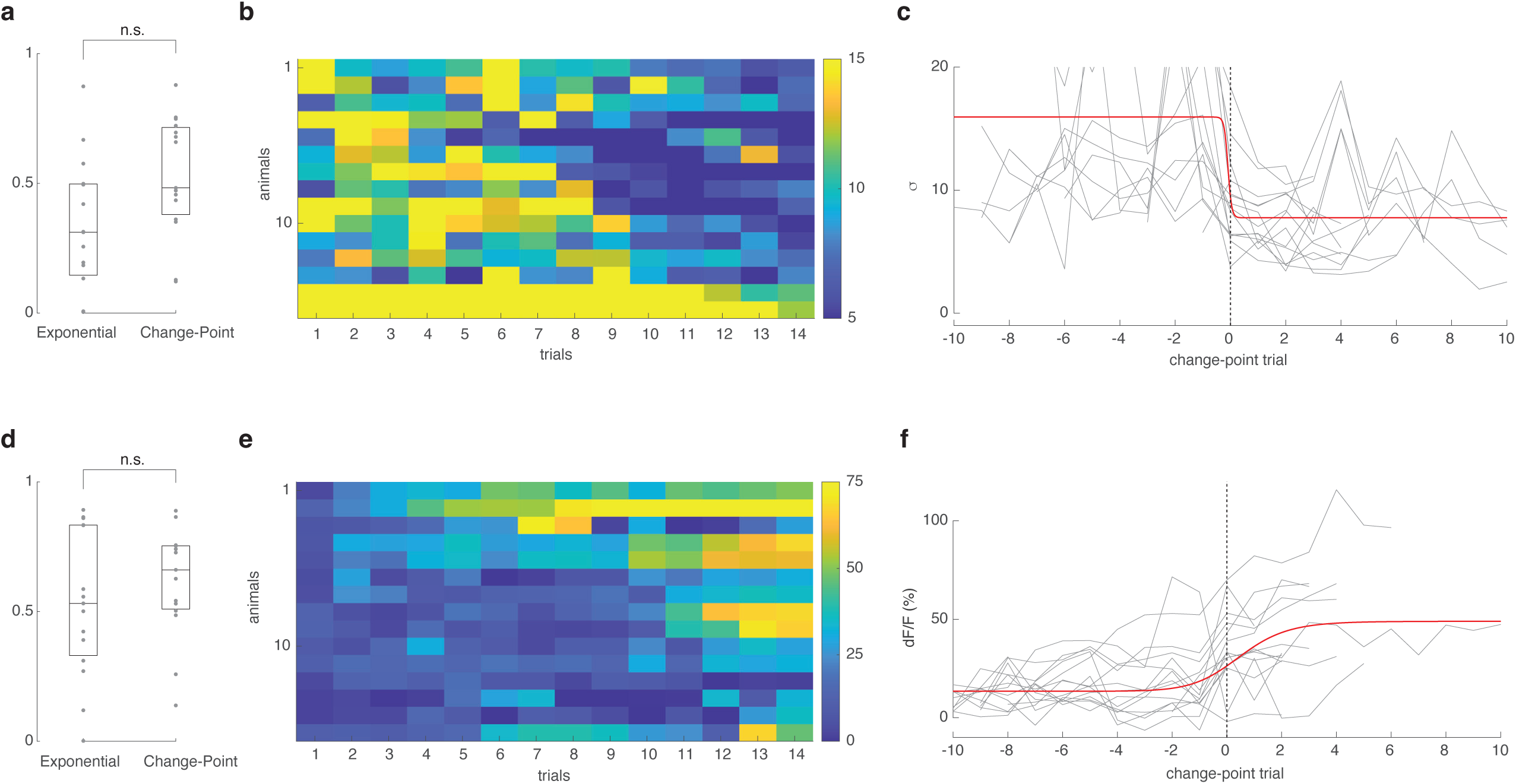
EB-PPM3 calcium dynamics during trace conditioning reveals that increasing the trace interval to 10s between CS and US further delays state transition trials. **(a)** R-squared goodness of fit of exponential and change-point models applied to per-trial standard deviations of dF_ratio_/F_ratio_ perturbation variability (relative to pre-CS baseline mean) 5 s after US (35-60 s) during trace conditioning with a 10 s trace interval (n = 15 flies). **(b)** Heatmap of modeled data from **(a)** in ascending order of each animal’s change point trial that partitions the data into two regions, minimizing the sum of the residual (squared) error of each region from its local mean. **(c)** Same data as **(b)**, trial aligned to center (trial 0) on each animal’s respective change point trial (gray lines). Red curve indicates sigmoidal curve fit to the per-trial median of the change point trial-aligned data. **(d)** R-squared goodness of fit of exponential and change-point models applied to per-trial mean pre-CS baseline dF_ratio_/F_ratio_ activity (0-10 s) during trace conditioning with a 10 s trace interval (n = 15 flies). **(e)** Heatmap of modeled data from **(d)** in ascending order of each animal’s change point trial that partitions the data into two regions, minimizing the sum of the residual (squared) error of each region from its local mean. **(f)** Same data as **(e)**, trial aligned to center (trial 0) on each animal’s respective change point trial (gray lines). Red curve indicates sigmoidal curve fit to the per-trial median of the change point trial-aligned data. Boxplot center (median), and edges (IQR). Scatters represent single-fly metrics. Groups compared using unpaired two-sided Mann–Whitney U tests. n.s. indicates not significant, * is p-value < 0.05, ** p-value < 0.01.

## References

1. Montague, P. R., Dayan, P. & Sejnowski, T. J. A framework for mesencephalic dopamine systems based on predictive Hebbian learning. J. Neurosci. 16, 1936–1947 (1996).

2. Sutton, R. S. & Barto, A. G. Reinforcement Learning: An Introduction.

3. Schultz, W., Dayan, P. & Montague, P. R. A Neural Substrate of Prediction and Reward. Science 275, 1593–1599 (1997).

4. Hollerman, J. R. & Schultz, W. Dopamine neurons report an error in the temporal prediction of reward during learning. Nat Neurosci 1, 304–309 (1998).

5. Schultz, W. Dopamine reward prediction error coding. Dialogues in Clinical Neuroscience 18, 23–32 (2016).

6. Schultz, W. Updating dopamine reward signals. Current Opinion in Neurobiology 23, 229–238 (2013).

7. Bayer, H. M. & Glimcher, P. W. Midbrain Dopamine Neurons Encode a Quantitative Reward Prediction Error Signal. Neuron 47, 129–141 (2005).

8. Gershman, S. J. & Niv, Y. Learning latent structure: carving nature at its joints. Current Opinion in Neurobiology 20, 251–256 (2010).

9. Gershman, S. J., Blei, D. M. & Niv, Y. Context, learning, and extinction. Psychological Review 117, 197–209 (2010).

10. Niv, Y. Learning task-state representations. Nat Neurosci 22, 1544–1553 (2019).

11. Courville, A. C., Daw, N. D. & Touretzky, D. S. Bayesian theories of conditioning in a changing world. Trends Cogn Sci 10, 294–300 (2006).

12. Nassar, M. R., Wilson, R. C., Heasly, B. & Gold, J. I. An Approximately Bayesian Delta-Rule Model Explains the Dynamics of Belief Updating in a Changing Environment. J. Neurosci. 30, 12366–12378 (2010).

13. Starkweather, C. K., Babayan, B. M., Uchida, N. & Gershman, S. J. Dopamine reward prediction errors reflect hidden-state inference across time. Nat Neurosci 20, 581–589 (2017).

14. Babayan, B. M., Uchida, N. & Gershman, S. J. Belief state representation in the dopamine system. Nat Commun 9, 1891 (2018).

15. Gershman, S. J. Dopamine, Inference, and Uncertainty. Neural Computation 29, 3311–3326 (2017).

16. Starkweather, C. K., Gershman, S. J. & Uchida, N. The Medial Prefrontal Cortex Shapes Dopamine Reward Prediction Errors under State Uncertainty. Neuron 98, 616–629.e6 (2018).

17. Waddell, S. Reinforcement signalling in Drosophila; dopamine does it all after all. Curr Opin Neurobiol 23, 324–329 (2013).

18. Aso, Y. et al. The neuronal architecture of the mushroom body provides a logic for associative learning. eLife 3, e04577 (2014).

19. Mushroom body output neurons encode valence and guide memory-based action selection in Drosophila | eLife. https://elifesciences.org/articles/04580.

20. Hige, T., Aso, Y., Modi, M. N., Rubin, G. M. & Turner, G. C. Heterosynaptic Plasticity Underlies Aversive Olfactory Learning in Drosophila. Neuron 88, 985–998 (2015).

21. Srinivasan, S., Greenspan, R. J., Stevens, C. F. & Grover, D. Deep(er) Learning. J. Neurosci. 38, 7365–7374 (2018).

22. Séjourné, J. et al. Mushroom body efferent neurons responsible for aversive olfactory memory retrieval in Drosophila. Nat Neurosci 14, 903–910 (2011).

23. Owald, D. et al. Activity of defined mushroom body output neurons underlies learned olfactory behavior in Drosophila. Neuron 86, 417–427 (2015).

24. Adel, M. & Griffith, L. C. The Role of Dopamine in Associative Learning in Drosophila: An Updated Unified Model. Neurosci Bull 37, 831–852 (2021).

25. Bennett, J. E. M., Philippides, A. & Nowotny, T. Learning with reinforcement prediction errors in a model of the Drosophila mushroom body. Nat Commun 12, 2569 (2021).

26. Jürgensen, A.-M., Sakagiannis, P., Schleyer, M., Gerber, B. & Nawrot, M. P. Prediction error drives associative learning and conditioned behavior in a spiking model of Drosophila larva. iScience 27, 108640 (2024).

27. Grover, D. et al. Differential mechanisms underlie trace and delay conditioning in Drosophila. Nature 603, 302–308 (2022).

28. Fisher, Y. E., Marquis, M., D’Alessandro, I. & Wilson, R. I. Dopamine promotes head direction plasticity during orienting movements. Nature 612, 316–322 (2022).

29. Bayer, H. M. & Glimcher, P. W. Midbrain dopamine neurons encode a quantitative reward prediction error signal. Neuron 47, 129–141 (2005).

30. Gallistel, C. R., Fairhurst, S. & Balsam, P. The learning curve: Implications of a quantitative analysis. Proceedings of the National Academy of Sciences 101, 13124–13131 (2004).

31. Durstewitz, D., Vittoz, N. M., Floresco, S. B. & Seamans, J. K. Abrupt transitions between prefrontal neural ensemble states accompany behavioral transitions during rule learning. Neuron 66, 438–448 (2010).

32. Rescorla, R. & Wagner, A. A theory of Pavlovian conditioning: Variations in the effectiveness of reinforcement and nonreinforcement. in Classical Conditioning II: Current Research and Theory vol. Vol. 2 (1972).

33. Gershman, S. J., Blei, D. M. & Niv, Y. Context, learning, and extinction. Psychol Rev 117, 197–209 (2010).

34. Rabiner, L. R. A tutorial on hidden Markov models and selected applications in speech recognition. Proceedings of the IEEE 77, 257–286 (1989).

35. Kemere, C. et al. Detecting Neural-State Transitions Using Hidden Markov Models for Motor Cortical Prostheses. Journal of Neurophysiology 100, 2441–2452 (2008).

36. Escola, S., Fontanini, A., Katz, D. & Paninski, L. Hidden Markov models for the stimulus-response relationships of multistate neural systems. Neural Comput 23, 1071–1132 (2011).

37. Soares, S., Atallah, B. V. & Paton, J. J. Midbrain dopamine neurons control judgment of time. Science 354, 1273–1277 (2016).

38. Mikhael, J. G. & Gershman, S. J. Adapting the flow of time with dopamine. J Neurophysiol 121, 1748–1760 (2019).

39. Dill, M., Wolf, R. & Heisenberg, M. Visual pattern recognition in Drosophila involves retinotopic matching. Nature 365, 751–753 (1993).

40. Gershman, S. J. & Uchida, N. Believing in dopamine. Nat Rev Neurosci 20, 703–714 (2019).

41. Langdon, A. J., Sharpe, M. J., Schoenbaum, G. & Niv, Y. Model-based predictions for dopamine. Current Opinion in Neurobiology 49, 1–7 (2018).

42. Langdon, A. J., Song, M. & Niv, Y. Uncovering the ‘state’: tracing the hidden state representations that structure learning and decision-making. Behav Processes 167, 103891 (2019).

43. Mermillod, M., Bugaiska, A. & Bonin, P. The stability-plasticity dilemma: investigating the continuum from catastrophic forgetting to age-limited learning effects. Front. Psychol. 4, (2013).

44. Behrens, T. E. J., Woolrich, M. W., Walton, M. E. & Rushworth, M. F. S. Learning the value of information in an uncertain world. Nat Neurosci 10, 1214–1221 (2007).

45. Nassar, M. R. & Troiani, V. The stability flexibility tradeoff and the dark side of detail. Cogn Affect Behav Neurosci 21, 607–623 (2021).

46. MacDonald, C. J., Lepage, K. Q., Eden, U. T. & Eichenbaum, H. Hippocampal “Time Cells” Bridge the Gap in Memory for Discontiguous Events. Neuron 71, 737–749 (2011).

47. Kropf, J., Talbot, C. B. & Miesenböck, G. A neuronal population clock for interval timing in Drosophila. Current Biology 36, 835–845.e8 (2026).

48. Hamou, N., Gershman, S. J. & Reddy, G. Reconciling time and prediction error theories of associative learning. Nat Commun 16, 10265 (2025).

49. Srinivasan, S. et al. Effects of stochastic coding on olfactory discrimination in flies and mice. PLOS Biology 21, e3002206 (2023).

50. Hige, T., Aso, Y., Modi, M. N., Rubin, G. M. & Turner, G. C. Heterosynaptic Plasticity Underlies Aversive Olfactory Learning in *Drosophila*. Neuron 88, 985–998 (2015).

51. Gibbon, J. Scalar expectancy theory and Weber’s law in animal timing. Psychological Review 84, 279–325 (1977).

52. Weir, P. T. et al. Anatomical Reconstruction and Functional Imaging Reveal an Ordered Array of Skylight Polarization Detectors in Drosophila. J. Neurosci. 36, 5397–5404 (2016).

53. Aso, Y. et al. The neuronal architecture of the mushroom body provides a logic for associative learning. eLife 3, e04577 (2014).

